# Transcriptomics reveal stretched human pluripotent stem cell-derived cardiomyocytes as an advantageous hypertrophy model

**DOI:** 10.1101/2021.12.13.472451

**Authors:** Lotta Pohjolainen, Heikki Ruskoaho, Virpi Talman

**Affiliations:** Drug Research Program and Division of Pharmacology and Pharmacotherapy, Faculty of Pharmacy, University of Helsinki, FI-00014 Helsinki, Finland

**Keywords:** Cardiomyocytes, Induced Pluripotent Stem Cells, Gene Expression, Transcriptomics, Left Ventricular Hypertrophy

## Abstract

Left ventricular hypertrophy, characterized by hypertrophy of individual cardiomyocytes, is an adaptive response to an increased cardiac workload that eventually leads to heart failure. Previous studies using neonatal rat ventricular myocytes (NRVMs) and animal models have revealed several genes and signaling pathways associated with hypertrophy and mechanical load. However, these models are not directly applicable to humans. Here, we studied the effect of cyclic mechanical stretch on gene expression of human induced pluripotent stem cell-derived cardiomyocytes (hiPSC-CMs) using RNA sequencing. hiPSC-CMs showed distinct hypertrophic changes in gene expression at the level of individual genes and in biological processes. We also identified several differentially expressed genes that have not been previously associated with cardiomyocyte hypertrophy and thus serve as attractive targets for future studies. When compared to previously published data attained from stretched NRVMs and human embryonic stem cell-derived cardiomyocytes, hiPSC-CMs displayed a smaller number of changes in gene expression, but the differentially expressed genes revealed more pronounced enrichment of hypertrophy-related biological processes and pathways. Overall, these results establish hiPSC-CMs as a valuable *in vitro* model for studying human cardiomyocyte hypertrophy.

**Non-standard Abbreviations and Acronyms:** ET-1, endothelin-1; GO, gene ontology; hESC-CM, human embryonic stem cell-derived cardiomyocyte; hiPSC, human induced pluripotent stem cell; hiPSC-CM, human induced pluripotent stem cell-derived cardiomyocyte; MAPK, mitogen-activated protein kinase; MEK1/2, mitogen-activated protein kinase kinase 1/2; NRVM, neonatal rat ventricular myocyte; PKC, protein kinase C; PBS, phosphate-buffered saline; RB+, RPMI 1640 supplemented with B-27; RB-, RPMI 1640 medium supplemented with B-27 without insulin; RT, room temperature; TF, transcription factor

## Introduction

The prevalence of cardiovascular diseases, including coronary artery disease and hypertension, is increasing rapidly, from approximately 271 million in 1990 to 523 million in 2019.^1^ However, treatment strategies have not evolved correspondingly; hence, cardiovascular diseases are the leading cause of death.^2^ Hypertension and myocardial infarction increase cardiac workload, causing structural and functional changes in the myocardium.^3^ These changes include left ventricular hypertrophy, which is characterized by cardiomyocyte enlargement. Although it is initially an adaptive response to physiological and pathological stimuli, such as mechanical stretch or neurohumoral activation, prolonged hypertrophy leads to contractile dysfunction and heart failure.

In response to hypertrophic stimuli, cardiomyocytes not only increase in size but also increase their protein synthesis, sarcomeres become disorganized, and specific changes in gene expression occur.^4^ The early genetic response to stretch is activation of immediate early response genes, such as proto-oncogenes *FOS* and *JUN*, components of the transcription factor AP-1. This is followed by upregulation of natriuretic peptide B (BNP) coding gene (*NPPB*) and reactivation of fetal genes such as natriuretic peptide A (*NPPA*), myosin-7 (*MYH7*), and skeletal muscle α-actin (*ACTA1*). A variety of effectors and signaling pathways mediate the hypertrophic response.^5^ For example, studies in animal models and in neonatal rat ventricular myocytes (NRVMs) have identified mitogen-activated protein kinase (MAPK) and protein kinase C (PKC) as potential mediators transducing the hypertrophic response in cardiomyocytes.^6,7^ However, controversial results of the exact role of these signal transducers have also been published, which can partially be explained by the use of different experimental models.

Cardiomyocyte hypertrophy has commonly been studied in animal models *in vivo* or in isolated NRVMs *in vitro*.^5^ However, studies conducted with animals or animal cells are not always directly translatable to humans. Human induced pluripotent stem cell-derived cardiomyocytes (hiPSC-CMs) offer a unique possibility to investigate human cardiomyocytes and abolish the effects of species differences. However, although hiPSC-CMs beat spontaneously, they are relatively immature by structural, metabolic and electrophysiological properties.^8,9^ We and others have previously shown that hiPSC-CMs respond to endothelin-1 (ET-1) by increasing the expression of pro-B-type natriuretic peptide (proBNP) and the corresponding gene (*NPPB*) along with other hypertrophy-related genes, although no morphological change was observed.^10,11^ On the other hand, Földes et al. showed that hiPSC-CMs lack the hypertrophic response to α-adrenergic stimuli.^9^ Hence, it seems that not all hypertrophic signaling pathways are functional in hiPSC-CMs.

The aim of this study was to characterize the transcriptomic response of hiPSC-CMs to mechanical stretch. In addition, we compared stretch-induced gene expression changes to previously published data from stretched NRVMs and human embryonic stem cell-derived cardiomyocytes (hESC-CMs).^12,13^ We also utilized our model to test the involvement of selected signaling pathways in mechanical load-induced hypertrophy of hiPSC-CMs by using pharmacological inhibition.

## Methods

### Compounds and reagents

The mitogen-activated protein kinase kinase 1/2 (MEK1/2) inhibitor U0126 and the p38 MAPK inhibitor SB203580 were purchased from Tocris Bioscience (Bristol, UK), the pan-PKC inhibitor Gö6983 was purchased from STEMCELL Technologies (Vancouver, Canada) and the classical PKC inhibitor Gö6976 was purchased from Merck (Darmstadt, Germany). Growth factor-reduced Matrigel was purchased from Corning (Bedford, MA, USA), and the small-molecule inhibitors Y-27632, CHIR99021 and Wnt-C59 were purchased from Tocris Bioscience (Bristol, UK). All other cell culture reagents were purchased from Gibco (Paisley, UK).

### Human induced pluripotent stem cell-derived cardiomyocytes

The hiPS(IMR90)-4 line was purchased from WiCell (Madison, WI, USA). Cells were maintained in Essential 8 medium on Matrigel-coated 6-well plates at 37 °C in a humidified atmosphere of 5% CO_2_. Cells were passaged 1:15 approximately every four days using Versene and regularly tested negative for mycoplasma contamination. Cardiomyocytes were differentiated as described previously.^10,14,15^ When the cultures were 80-95% confluent, differentiation was initiated by the addition of 6 µM CHIR99021 (Day 0) to RPMI 1640 medium supplemented with B-27 without insulin (RB-). On Day 1 or Day 2, the medium was changed to fresh RB-. On Day 3, fresh RB-containing 2.5 µM Wnt-C59 was added. On Days 5, 7 and 9, RB-was changed to fresh RB-. On Day 11, metabolic selection of cardiomyocytes was initiated by changing the RB-to RPMI 1640 without glucose supplemented with B-27 (with insulin). On Day 13, the cells were fed fresh metabolic selection medium. From Day 15 onwards, the cardiomyocytes were cultured in RPMI 1640 supplemented with B-27 (RB+). On Day 15 or 17, the differentiated hiPSC-CMs (≥ 95% pure cardiomyocyte cultures) were dissociated and seeded at a density of 700,000-800,000 cells/well in RB+ supplemented with 10% fetal bovine serum (FBS) on flexible collagen I-coated 6-well BioFlex® culture plates (Flexcell International Corporation, Hillsborough, NC, USA) with additional Matrigel coating. The hiPSC-CMs were allowed to attach for 48 h, after which the medium was changed to serum-free RB+. Cardiomyocytes were maintained by changing fresh RB+ approximately every four days until the start of experiments on Days 29-43.

### Cyclic mechanical stretch

The hiPSC-CMs were exposed to cyclic mechanical stretch by applying vacuum suction to the BioFlex® plates with an FX-5000 Tension System (Flexcell International Corporation). For pharmacological assays, compounds or vehicle (0.1% DMSO) were added one hour before starting the stretch. Equibiaxial stretch was applied for 24 h, 48 h or 72 h in two-second cycles (0.5 Hz) to induce cyclic stretch varying from 10% to 21% elongation, corresponding to 42-80 kPa pressure (Figure S1). Unstretched control cells were from the same differentiation and were seeded on BioFlex® plates at the same time, but no stretch was applied. The schematic of the experimental design is shown in Figure 1.

**Figure 1.**
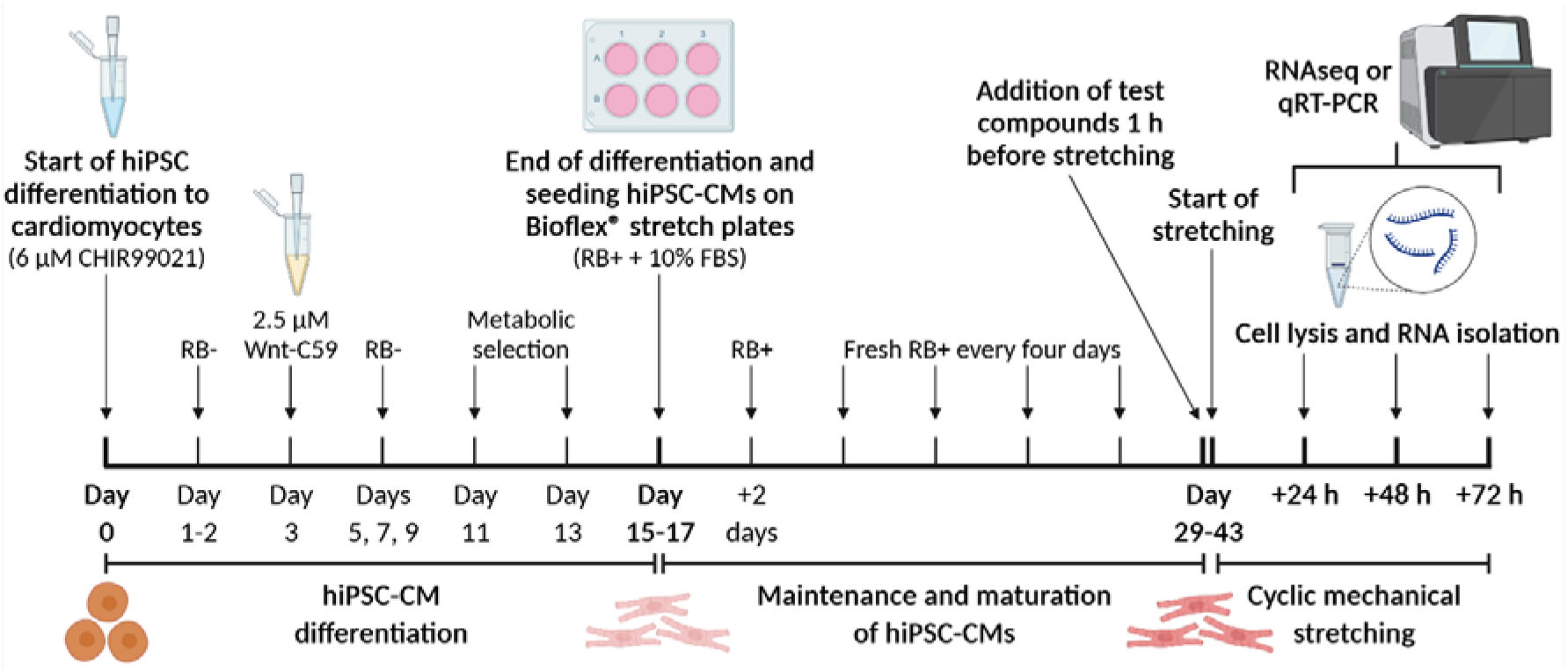
Schematic representation of the experimental design. Human induced pluripotent stem cells (hiPSCs) were differentiated to cardiomyocytes (hiPSC-CMs) using small molecules: GSK3 inhibitor CHIR99021 and Wnt pathway inhibitor C59. Metabolic selection by glucose deprivation resulted ≥ 95% pure cardiomyocyte cultures. After 2-4 weeks of maintenance and maturation, hiPSC-CMs were exposed to test compounds and cyclic mechanical stretch, after which hiPSC-CMs were lyzed and RNA was analyzed. RB-, RPMI 1640 medium supplemented with B-27 without insulin; RB+, RPMI 1640 supplemented with B-27; FBS, fetal bovine serum. Figure was created with BioRender.com.

### RNA isolation

Cells were lysed in 350 µl of RA1 lysis buffer (Macherey-Nagel, Düren, Germany) supplemented with 1% β-mercaptoethanol and stored at -80 °C (maximum one month) before RNA isolation. RNA was isolated using a NucleoSpin RNA kit (Macherey-Nagel) according to the manufacturer’s instructions. Analysis of the RNA concentration and quality was performed with a NanoDrop 1000 spectrophotometer (Thermo Fisher Scientific, Waltham, MA, USA) for qRT-PCR and with 4200 TapeStation (Agilent, Santa Clara, CA, USA) for RNA sequencing (RNAseq). One of the sample pairs at 24 h time point was omitted from the RNAseq due to poor quality (RNA integrity number < 9).

### Quantitative Reverse Transcription PCR (qRT-PCR)

cDNA was synthesized from 100–500 ng of total RNA in 10 µl reactions with a Transcriptor First Strand cDNA Synthesis Kit (Roche, Basel, Switzerland) according to the manufacturer’s protocol using random hexamer primers and an MJ Mini Personal thermal cycler (Bio-Rad, Hercules, CA, USA). The cDNA was diluted 1:10 in PCR grade H_2_O and stored at -20 °C. Commercial TaqMan® Gene Expression Assays (Thermo Fisher Scientific) listed in Table S1 were used with LightCycler® 480 Probes Master reagent (Roche) according to the manufacturer’s instructions. A LightCycler® 480 Real-Time PCR System (Roche) was used to analyze 4.5 µl of the cDNA dilution in 10 µl reactions on a white LightCycler® 480 Multiwell Plate 384 (Roche). To confirm the absence of PCR contamination, no-template controls were used. Each reaction was run at least in triplicate, and the average of the technical replicates was used in the analysis as n=1. Grubbs’ test was used to identify outliers within technical replicates and identified outliers were excluded from the analysis. The 2^-ΔΔCt^ method was used to analyze the relative gene expression using *ACTB* and *18S rRNA* as reference genes.

### RNA sequencing

RNAseq was performed as single-end sequencing for a read length of 75 bp with an Illumina NextSeq 500 sequencer (Illumina, San Diego, CA, USA) in high output runs using a NEBNext Ultra Directional RNA Library Prep kit (New England Biolabs, Ipswich, MA, USA) including rRNA depletion. Data quality was analyzed by FastQC, and quality trimming was applied to the data with Trimmomatic software.^16^ The sample reads were aligned against the Genome Reference Consortium Human Build 38 patch release 13 (GRCh38.p13, GCA_000001405.28) reference with Spliced Transcripts Alignment to a Reference (STAR).^17^ The mapping quality was assessed with Qualimap. Read quantification was created with featureCounts, and differential expression with quality assessment was performed with DESeq2.^18–20^

### Functional enrichment analysis of gene sets

Gene Ontology (GO) enrichment of differentially expressed genes was analyzed with GOrilla (version updated 27^th^ of February 2021, available at http://cbl-gorilla.cs.technion.ac.il/).^21^ The tool searched for GO terms that were enriched in the target set compared to the background set. A list of upregulated or downregulated genes was entered as target sets. As the background set, we used all expressed genes in our dataset (30,861 genes), defined as genes with a detected signal in at least two samples in one treatment group. A relatively low p value threshold of p<0.0001 was used for running the analysis, as no correction for multiple testing was applied. False discovery rate (FDR)-adjusted p values were calculated after the analysis, and an FDR-adjusted p value <0.05 was considered significant.

WebGestalt (version 2019, available at http://www.webgestalt.org/) was used for KEGG and Reactome pathway analyses and for transcription factor target analysis.^22^ The same target and reference gene sets as in the GO enrichment analysis were used. The Benjamini-Hochberg method was used for multiple testing, and a significance level of <0.05 was used. KEGG Mapper (available at https://www.genome.jp/kegg/mapper.html) was used to determine the cellular functions of proteins representing the differentially expressed genes.^23^ Encyclopedia of RNA Interactomes (ENCORI; available at http://starbase.sysu.edu.cn/) was used to predict the putative interaction partners of differentially expressed lncRNAs.^24^

### Dataset comparison

Our dataset of stretched hiPSC-CMs was compared to datasets of the two other studies. We used data from differentially expressed genes of stretched NRVMs by Rysä et al., who isolated NRVMs from 2-to 4-day-old Sprague–Dawley rats and stretched cells with the FlexCell vacuum system, similar to the present study (0.5 Hz, 10-25% elongation).^12^ We also compared our data to the data of stretched hESC-CMs obtained by Ovchinnikova et al., who had used a slightly different stretching protocol: cyclic stretch with elongation from 0% to 15% was applied at a frequency of 1 Hz with the FlexCell system.^13^

### Immunofluorescence staining, imaging and image analysis

Brefeldin A (Invitrogen, Carlsbad, CA, USA) was added to the cells 3 h before fixation to block exocytosis of proBNP-containing vesicles. At the end of stretching, the cells were washed twice with phosphate-buffered saline (PBS) and fixed with 4% paraformaldehyde at room temperature (RT) for 15 min followed by 3×5 min washes with PBS. A square piece (approximately 10 mm x 10 mm) was cut from the center of each flexible bottomed well of Bioflex® plate for staining. Subsequently, the cells were permeabilized with 0.1% Triton X-100 in PBS at RT for 10 min. Non-specific binding sites were blocked with 4% FBS in PBS for 45 min followed by addition of primary antibodies diluted in 4% FBS in PBS. Cardiac troponin T (cTnT) antibody (ab45932, Abcam) was used at 1:800 and proBNP antibody (ab13115, Abcam) at 1:200. For the secondary antibody staining control, primary antibodies were omitted. After a 60-min incubation at RT, the cells were washed 3×5 min with PBS and incubated with Alexa Fluor® -conjugated secondary antibodies (Life Technologies, Eugene, Oregon) at 1:200, and 4′,6-diamidino-2-phenylindole (DAPI; Sigma-Aldrich) at 1 µg/ml at RT for 45 min. The cells were then washed 3×5 min with PBS and mounted between a microscope slide and a coverslip using ProLong™ Gold Antifade Mountant (Invitrogen). The samples were cured at RT overnight and then stored at 4°C until imaged. The cells were imaged with ImageXpress Micro Confocal imaging system (Molecular Devices) using Nikon 10x Plan Apo 0.5 NA air objective and 40x C CFI APO LWD Lambda S 1.15 NA water immersion objective. MetaXpress software (Molecular Devices) was used to analyze proBNP and cTnT intensity in cardiomyocyte nuclei and perinuclear area. First, background was subtracted from DAPI and proBNP images using the top hat filter. The nuclei were then identified based on DAPI staining and the cardiomyocyte cytoplasm was identified based on cTnT staining. The average proBNP staining intensity was quantified from the perinuclear region of cardiomyocytes defined as a 10-pixel ring around each cTnT positive nucleus. The average cTnT staining intensity was measured from combined nuclear and perinuclear area.

### Statistical analysis

In statistical analysis, each treatment group comprised 3–5 independent experiments of cells from individual differentiations. Sample size was determined by initial analysis of *NPPA* and *NPPB* responses measured by qRT-PCR and previous study on NRVMs.^12^ Each qRT-PCR included technical replicates, and the average was calculated for statistical analysis to represent n=1. Statistical analysis of the qRT-PCR results was performed in IBM SPSS Statistics 25 software. Student’s t-test for independent samples was performed to compare the stretched and unstretched samples. A nonparametric Mann–Whitney U test was applied for the normalized qRT-PCR data of the pharmacological inhibitor experiments. A value of p<0.05 was considered statistically significant. Statistical analysis of the RNAseq results was performed as a pairwise comparison of stretched and unstretched samples using the Wald test in DESeq2 software. Genes with a fold change (FC) >1.5 and Benjamini-Hochberg adjusted p<0.05 were defined as differentially expressed.

## Results

### Effects of mechanical stretch on natriuretic peptide expression in hiPSC-CMs

The mechanical stretch model of hiPSC-CMs (Figure 1) was first validated by measuring the mRNA expression of *NPPA* and *NPPB*, hallmark genes of cardiomyocyte hypertrophy.^25^ After 24 h of cyclic mechanical stretch, hiPSC-CMs showed increased expression of both *NPPA* and *NPPB* (Figures 2A and 2B). At 48 h and 72 h, the upregulation of the *NPPA* and *NPPB* mRNA levels was not statistically significant, although increased gene expression was observed in each independent experiment. To confirm the increased BNP expression in protein level, we stained proBNP (the N-terminal fragment of the BNP prohormone) in stretched hiPSC-CMs. An evident increase in perinuclear proBNP expression was observed after a 24-h stretch (Figure 2C and 2E). In agreement with the time-dependent gene expression, no apparent change in proBNP expression was observed after a 48-h stretch. However, there was a trend towards increased cTnT intensity with increasing duration of stretch (Figure 2D and 2E). In addition, high purity of cardiomyocytes was confirmed by staining for cTnT. High-resolution images acquired with 40x objective are available in Supplementary Material (Figure S2). No clear effect on sarcomere organization or alignment was observed.

**Figure 2.**
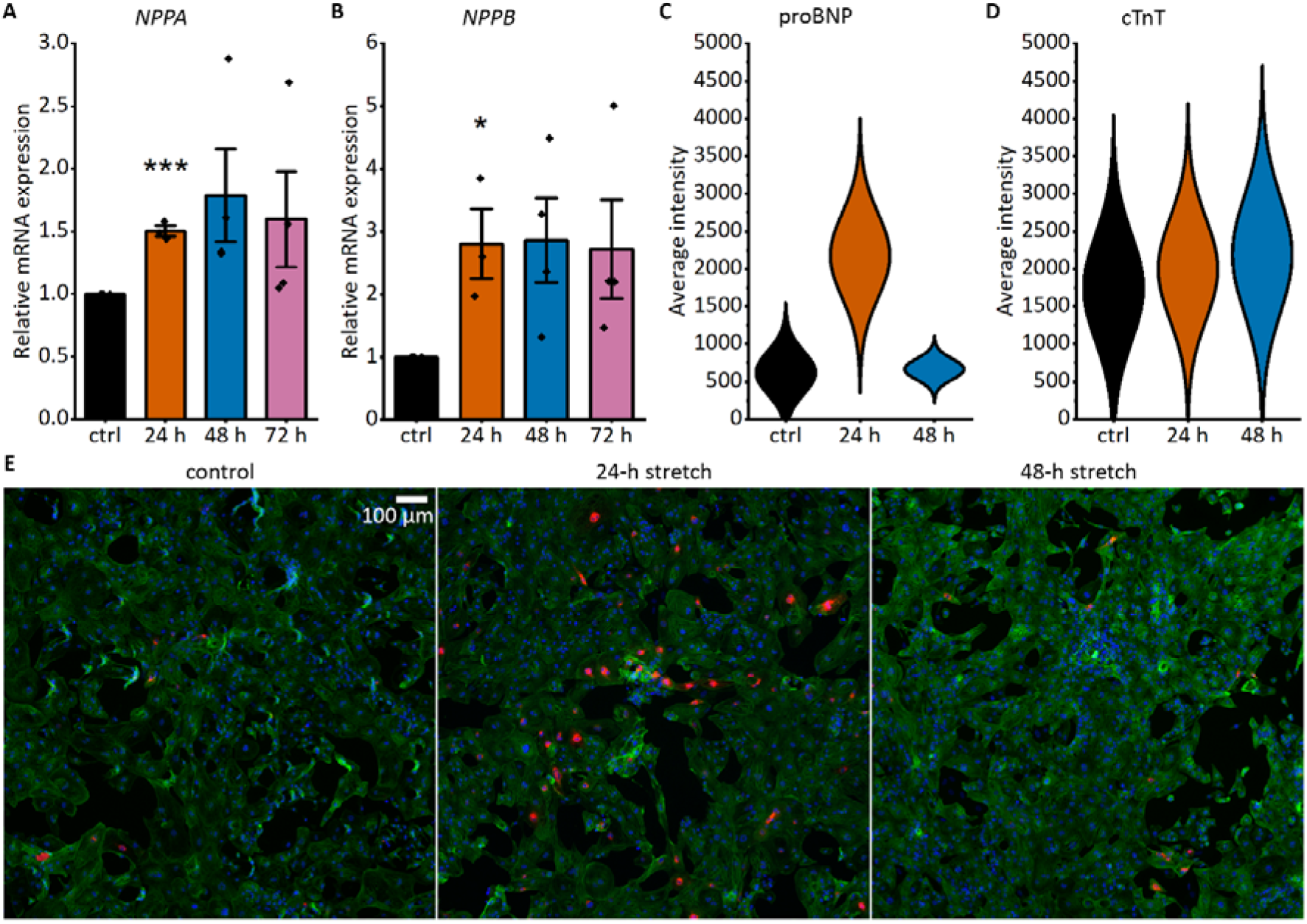
Cyclic mechanical stretching induces natriuretic peptide expression in human induced pluripotent stem cell-derived cardiomyocytes (hiPSC-CMs). A-B, hiPSC-CMs respond to stretch by increased expression of hypertrophy-associated genes *NPPA* (natriuretic peptide A; A) and *NPPB* (natriuretic peptide B; B). mRNA expression of 24-h, 48-h and 72-h stretched hiPSC-CMs measured with qRT-PCR was normalized to the unstretched control. The data are shown as mean ± standard error of the mean, and values from individual experiments are presented as dots (n=3-4, where n represents biological replicates from individual differentiations). *p<0.05, ***p<0.001 vs. unstretched control, Student’s t-test for independent samples. C, Average perinuclear proBNP intensity analyzed from 15,000 stained cardiomyocytes/sample, from a single differentiation. D, Average cardiac troponin T (cTnT) intensity in cardiomyocyte nuclear and perinuclear area analyzed from 15,000 stained cardiomyocytes/sample, from a single differentiation. E, Representative images of unstretched control, 24-h stretched, and 48-h stretched hiPSC-CMs acquired with 10x air objective. hiPSC-CMs were stained for DNA (DAPI; blue), pro-B-type natriuretic peptide (proBNP; red) and cTnT (green). Scale bar, 100 µm.

### Mechanical stretch-induced genome-wide gene expression program in hiPSC-CMs

To identify genome-wide gene expression changes regulated by mechanical stretch, we performed RNAseq from mechanically stretched hiPSC-CMs at 24 h, 48 h and 72 h and their unstretched controls. Principal component analysis using the whole gene set showed separation of individual experiments defined by the first principal component and separation of stretched and control samples at 24 h and 48 h defined by the second principal component (Figure S3). However, after 72 h of stretching, no clear difference between stretched and unstretched groups was detected, while more pronounced separation of individual experiments was seen instead. These findings suggest strong conserved early responses to stretch and increased biological variation over time.

Of the 30,861 genes identified in our samples, 134 genes showed differential expression (FC>1.5, FDR-adjusted p<0.05) after stretching. Our analysis identified 75, 28 and 2 upregulated genes in response to 24 h, 48 h and 72 h of stretch, respectively (Figure 3A). In addition, 12, 30 and 0 genes were downregulated in response to 24 h, 48 h and 72 h of stretch, respectively (Figure 3B). Venn diagrams demonstrated minor overlaps between time points, indicating time-dependent regulation of gene expression. The top 12 differentially expressed genes at each time point are presented in Figures 3C and 3D. The full data are available in Dataset S1.

**Figure 3.**
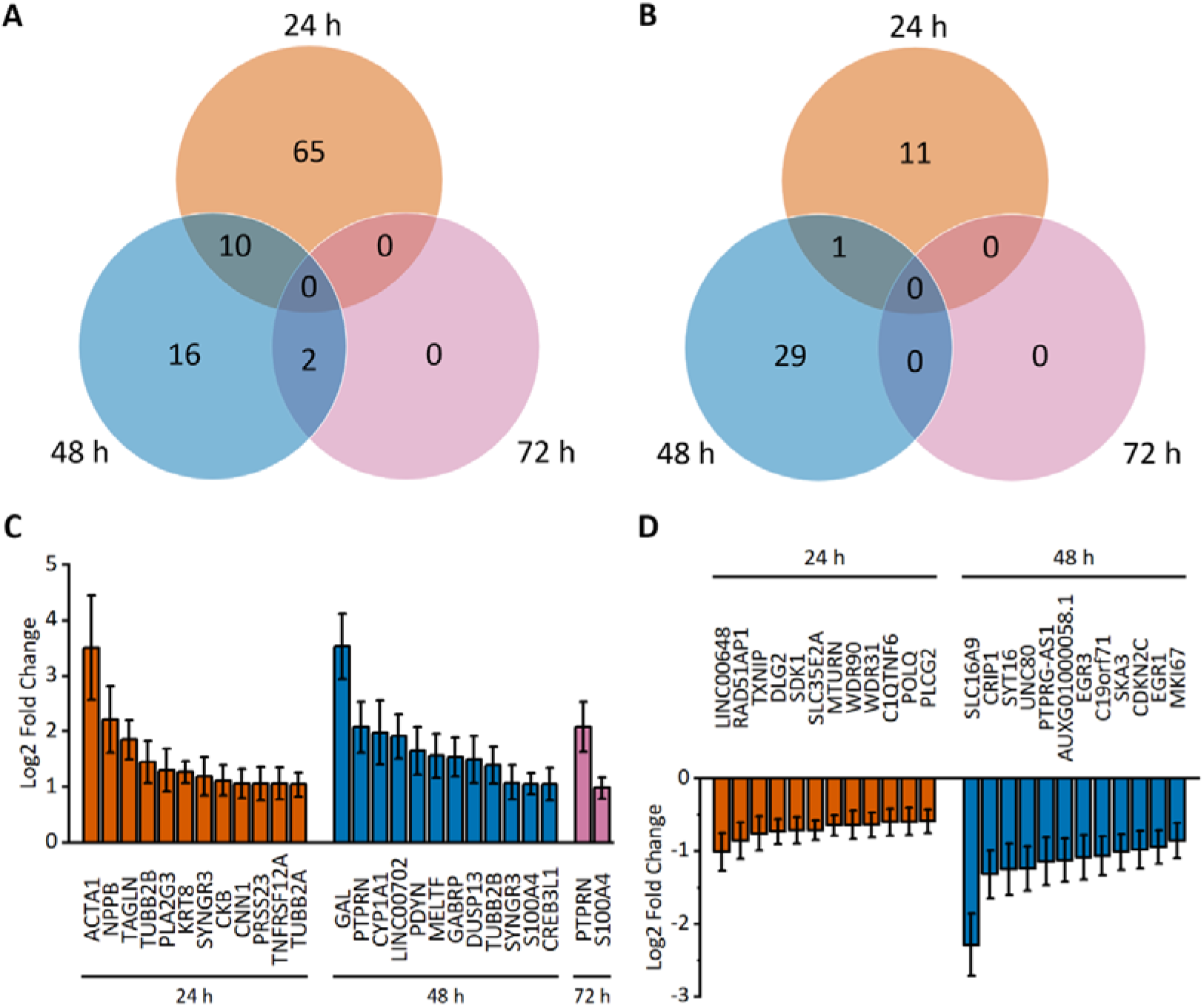
Mechanical stretch-induced differential gene expression of human induced pluripotent stem cell-derived cardiomyocytes (hiPSC-CMs). A-B, Venn diagrams show the number of upregulated (A) and downregulated (B) genes after 24 h, 48 h and 72 h of cyclic stretch measured with RNA sequencing. Differential expression was defined as a >1.5-fold change compared to the unstretched control. C-D, The expression of the top 12 up-(C) and downregulated (D) genes is presented as log2-fold change relative to the unstretched control ± standard error of the mean (n=3 for 24 h, n=4 for 48 h and 72 h; where n represents biological replicates of cells from individual differentiations). Only statistically significant (Benjamini-Hochberg adjusted p<0.05) results are presented.

Multiple hypertrophy-associated genes were upregulated. These include fetal genes coding for natriuretic peptides (*NPPA* and *NPPB*), skeletal muscle alpha actin (*ACTA1*), and transgelin (*TAGLN*), and several genes encoding contractile proteins, such as cardiac alpha actin (*ACTC1*), myosin light chain 3 (*MYL3*), troponins (*TNNI3, TNNC1*), and tropomyosin 2 (*TPM2)*. In addition to contractile proteins, other cytoskeletal proteins, such as alpha- and beta-tubulins (*TUBA4A, TUBA1A, TUBB2A, TUBB6*), alpha-actinin 1 (*ACTN1)*, nestin (*NES*) and keratins (*KRT8, KRT18*), were upregulated. These changes confirm that mechanical stretching of hiPSC-CMs induces alterations in gene expression, which are characteristic for cardiomyocyte hypertrophy. The upregulated genes mainly encode enzymes (26), exosomal proteins (20) and cytoskeletal proteins (15), while the downregulated genes encode enzymes (6) and transcription factors (6) (Table S2).

We selected eleven genes for validation by qRT-PCR. The selection was first based on differential expression of both up- and downregulated genes. Second, both protein-coding and noncoding (*LINC00648, PTPRG-AS1*) genes were selected. In addition, different protein-coding genes were selected, including hypertrophy-associated secreted peptides (*NPPA, NPPB*), cytoskeletal proteins (*ACTA1, ACTC1, ACTN1, TNNI3*), a transcription factor (*CSRP3*) and a transporter protein (*SLC16A9*). Overall, similar results were obtained with both qRT-PCR and RNAseq (Figure 4), except for *ACTN1*, which showed downregulation in qRT-PCR and slight upregulation in RNAseq after 72 h of stretch (Figure 4A).

**Figure 4.**
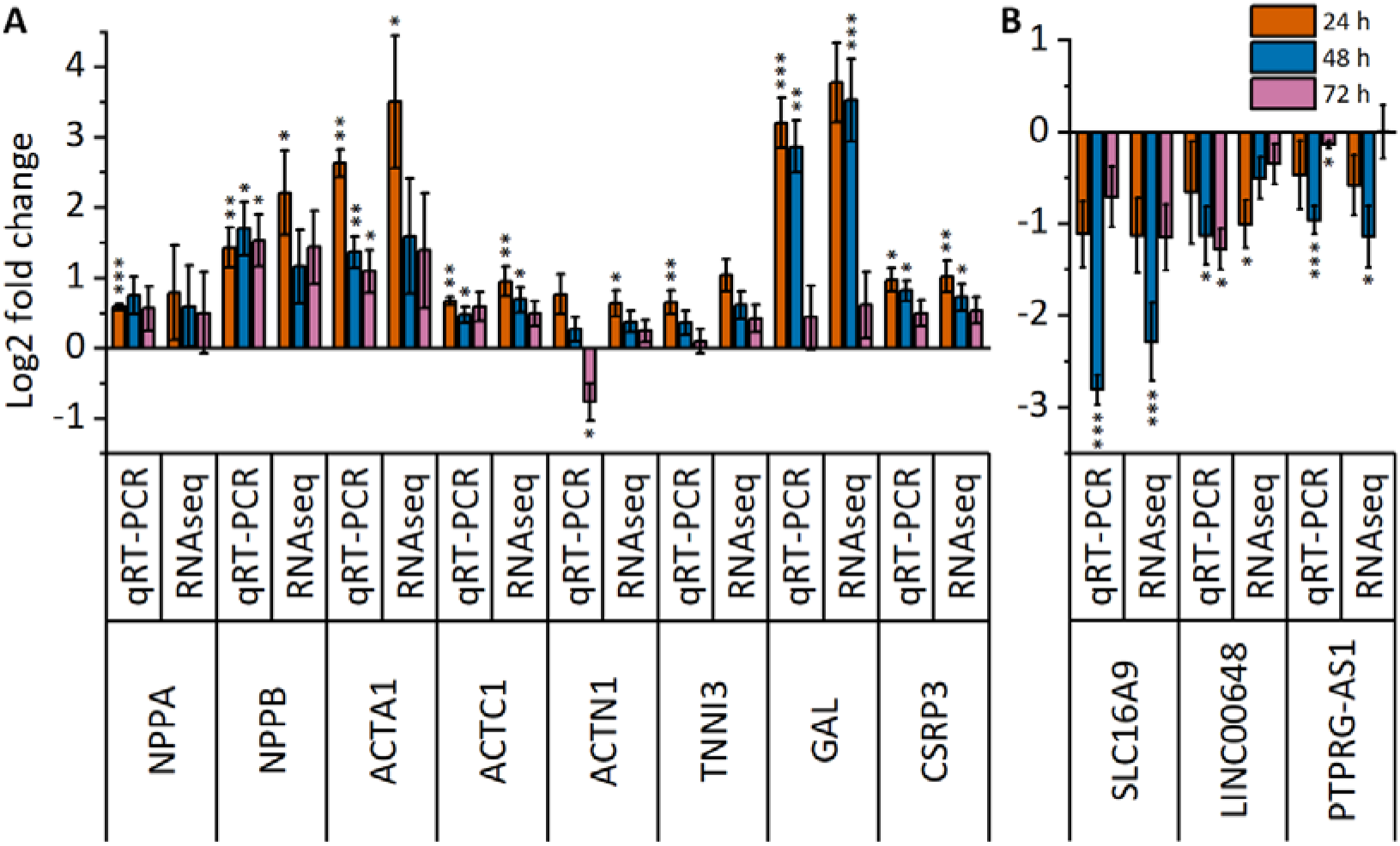
Time-dependent changes in gene expression of selected genes. Differential expression of 11 selected genes from RNA sequencing was validated by qRT-PCR after 24 h, 48 h and 72 h of stretch. (A) Upregulated genes; (B) downregulated genes. The results are presented as log2-fold change relative to the unstretched control ± standard error of the mean (n=3 for 24 h, n=4 for 48 h and 72 h; where n represents biological replicates of cells from individual differentiations). *p<0.05, **p<0.01, ***p<0.001 vs. unstretched control, Student’s t-test for independent samples (qRT–PCR), Wald test (RNAseq).

### Comparison of differentially expressed genes in hiPSC-CMs, NRVMs and hESC-CMs

We compared our data with the NRVM data published by Rysä et al. to elucidate similarities and differences between these two *in vitro* cardiomyocyte models from different species.^12^ Both we and Rysä et al. used time points of 24 h and 48 h; hence, these were selected for comparison, although the equivalency of these time points between the species has not been proven. Overall, the number of differentially expressed genes was drastically different; for example, after a 48-h stretch, over 600 genes were upregulated in NRVMs, while only 28 genes were upregulated in hiPSC-CMs (Figure 5). Interestingly, 21 differentially expressed genes showed similar changes in both cell models. In fact, 3 genes were upregulated in both cardiomyocyte types at both time points: *CASQ1, TIMP1* and *TUBB2B*. We did not identify genes that were consistently downregulated in both CM types at 24h and 48 h. After 24 h of stretch, 14 genes were upregulated in both cell types, while no commonly downregulated genes were identified (Figure 5C). Additionally, we identified 8 genes that were upregulated and 2 genes that were downregulated in both cell types after a 48-h stretch (Figure 5D). Cross comparison of different time points revealed 5 upregulated genes and 2 downregulated genes in hiPSC-CMs after 48 h of stretch with similar expression changes in NRVMs after 24 h of stretch (Table S3). In addition, 20 upregulated genes and one downregulated gene in hiPSC-CMs after a 24-h stretch were similarly differentially expressed in NRVMs after a 48-h stretch. Hence, the differentially expressed genes showed the most similarity between upregulated genes in hiPSC-CM at 24 h and in NRVMs at 48 h. Only three genes showed opposing expression in hiPSC-CMs and NRVMs: *MASP1, ENO3*, and *CES1* were upregulated in hiPSC-CMs but downregulated in NRVMs.

**Figure 5.**
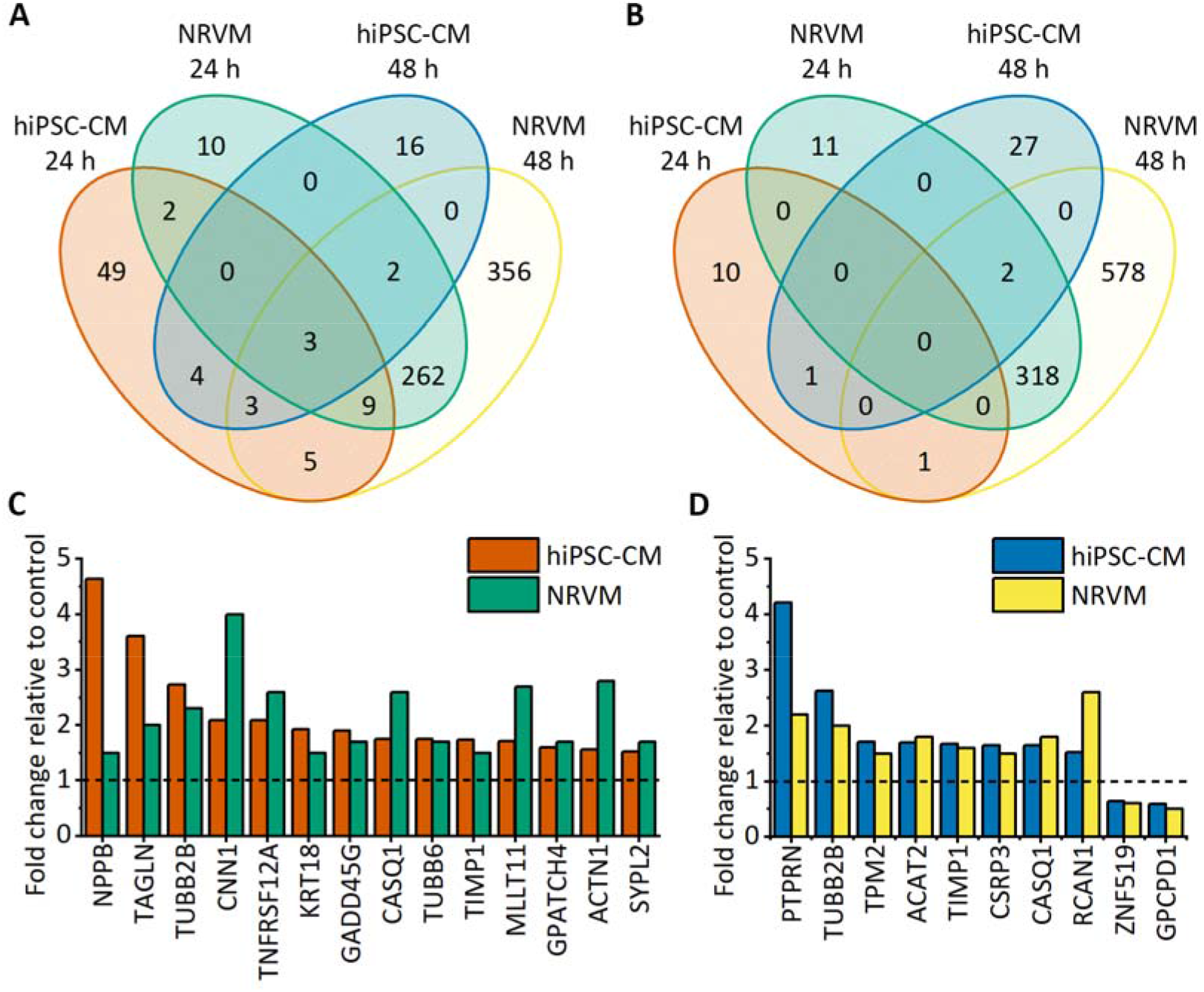
Comparison of differentially expressed genes in human induced pluripotent stem cell-derived cardiomyocytes (hiPSC-CMs) and neonatal rat ventricular myocytes (NRVMs) after 24 h and 48 h of stretch. NRVM gene expression data are from Rysä et al.^12^ A-B, Venn diagrams show the number of upregulated (A) and downregulated (B) genes after 24 h and 48 h of stretch in hiPSC-CMs and NRVMs. C-D, Expression of genes differentially regulated in both cell types after 24 h (C) and 48 h (D) of stretching normalized to the unstretched control (n=3 for 24 h hiPSC-CMs, n=4 for 48 h and 72 h hiPSC-CMs, and n=5 for NRVMs).

We also compared our differential gene expression data at 48 h with that from 48 h stretched hESC-CMs reported by Ovchinnikova et al.^13^ Of 936 differentially expressed genes in stretched hESC-CMs, only 13 genes were similarly expressed in hiPSC-CMs: four genes were upregulated, and nine genes were downregulated. The upregulated genes included *TUBB2B*, which was upregulated in all cell types (hiPSC-CMs, NRVMs and hESC-CMs) after 24 h (not studied in hESC-CMs) and 48 h of stretching. Other upregulated genes were *DUSP13, ACAT2* and *ENO3*. In addition, one of the genes downregulated in both hiPSC-CMs and NRVMs, *ZNF519*, was also downregulated in hESC-CMs. In hiPSC-CMs and hESC-CMs, no changes in opposite directions were observed in any differentially expressed gene. All common differentially expressed genes and their fold changes are presented in Table S3.

### Functional analysis of differentially expressed genes in stretched hiPSC-CMs

To identify the biological functions regulated by the differentially expressed genes, we performed GO enrichment analysis. The GOrilla (version updated 27^th^ of February 2021, available at http://cbl-gorilla.cs.technion.ac.il/) analysis recognized 18,992 genes out of 30,861 gene terms entered. Only 15,554 of these genes were associated with a GO term, and these were used for enrichment calculation. Enriched GO terms were found only for upregulated genes. The enriched biological processes of upregulated genes at all time points are presented in Figures 6A and S4. All enriched processes were highly related to cardiomyocyte hypertrophy. Actin filament-based movement, more specifically actin-myosin filament sliding and muscle filament sliding, was the most enriched biological process. Other enriched processes can be grouped under four categories: muscle contraction, secretion, regulation of cell death, and steroid biosynthesis. This result indicates that very specific biological processes are activated in hiPSC-CMs in response to stretching.

**Figure 6.**
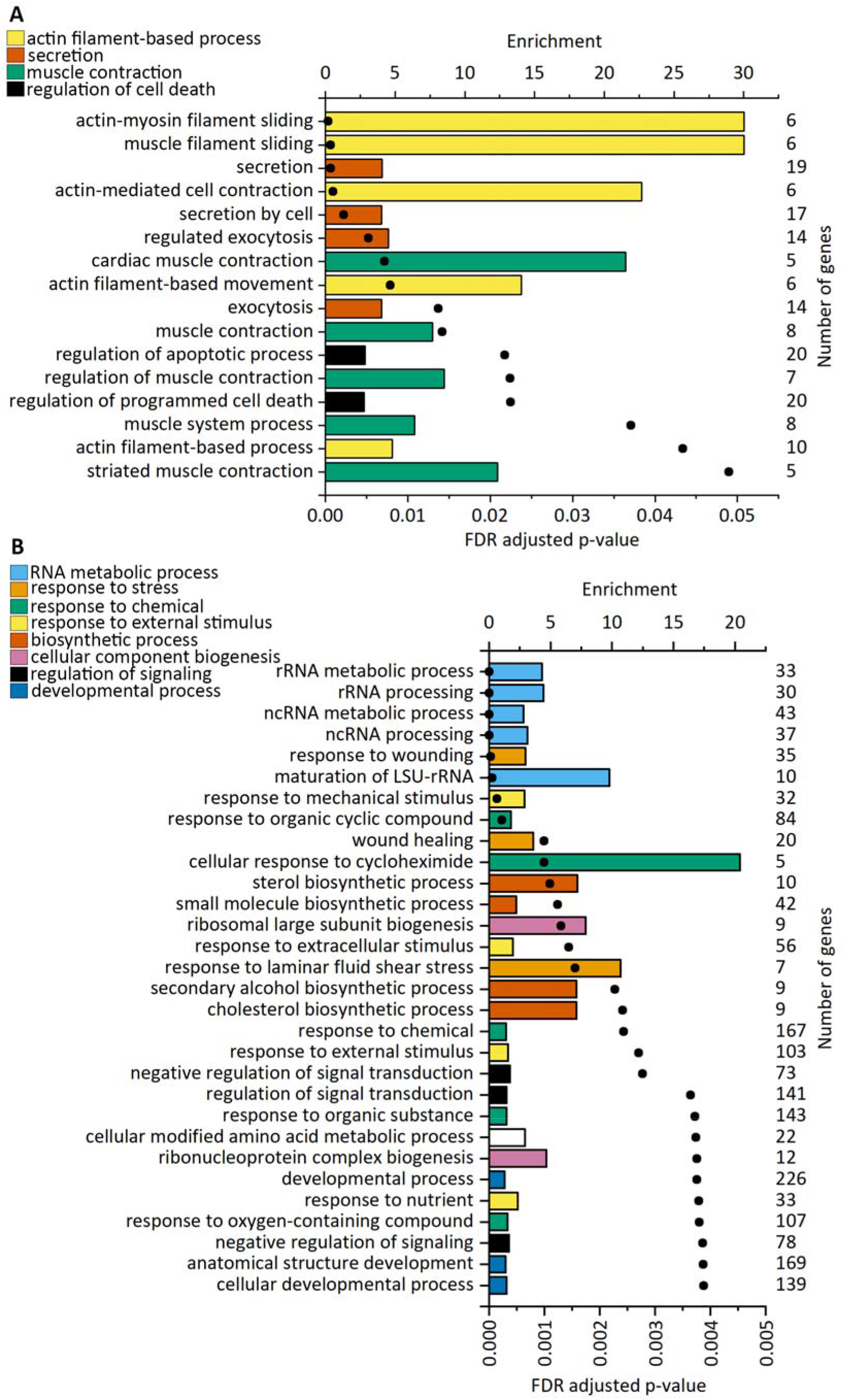
Enriched biological processes based on upregulated genes after cyclic stretch in human induced pluripotent stem cell-derived cardiomyocytes (hiPSC-CMs; A) and neonatal rat ventricular myocytes (NRVMs; B). Gene Ontology (GO) enrichment analysis was performed with GOrilla. For each significantly enriched GO term, enrichment values are presented as bars, and FDR-adjusted p values are presented as dots. The number of upregulated genes associated with each GO term is shown on the right. The upregulated genes in the NRVMs used for the analysis are from Rysä et al.^12^ The top 30 terms for NRVMs are shown.

We also searched for enriched GO terms for molecular function and cellular components. Again, enriched GO terms were only found for upregulated genes. Three molecular functions were significantly overrepresented: structural molecule activity, structural constituent of the cytoskeleton, and calcium-dependent protein binding (Figure S5). Among the cellular components, 14 GO terms, especially terms related to extracellular vesicles, supramolecular complexes and cytoskeleton, were enriched (Figure S5). These analyses confirm that structural and cytoskeletal protein-coding genes are among the most upregulated genes in stretched hiPSC-CMs. The complete data of the GO analyses are available in Dataset S2.

### Comparison of the functional analysis of differentially expressed genes in hiPSC-CMs and NRVMs

To compare stretch-induced enriched biological processes in hiPSC-CMs and NRVMs, we performed a similar GO analysis for upregulated genes in NRVMs reported by Rysä et al.^12^ In line with a higher number of upregulated genes in NRVMs compared to hiPSC-CMs, more enriched biological processes were found (71 GO terms). Hence, while a very limited number of specific processes were enriched in stretched hiPSC-CMs, a broad range of biological processes were detected in NRVMs. The most evidently enriched biological processes associated with upregulated genes in NRVMs were RNA metabolic processes, response to stimulus, biosynthetic processes, cellular component biogenesis, developmental processes, and regulation of cell death (Figure 6B). Upregulated genes from both hiPSC-CMs and NRVMs thus share GO terms associated with the regulation of apoptosis and steroid biosynthesis.

To further discover the functionality of the differentially expressed genes, KEGG and Reactome pathway analyses were performed. KEGG pathway analysis revealed 11 and 10 enriched pathways in upregulated genes in hiPSC-CMs and NRVMs, respectively (Figures 7A and 7B). However, none of the pathways was common to both cell types. In hiPSC-CM, the pathways included cardiac- and cardiomyocyte-associated pathways, while the enriched terms in NRVMs were heterogeneous, and half of them were cancer-associated. In turn, Reactome pathway analysis resulted in 31 and 17 enriched pathways for hiPSC-CMs and NRVMs, respectively (Figures 7C and 7D). Three pathways were enriched in both cell types: striated muscle contraction, HSP90 chaperone cycle for steroid hormone receptors (SHRs), and the role of GTSE1 in G2/M progression after the G2 checkpoint.

**Figure 7.**
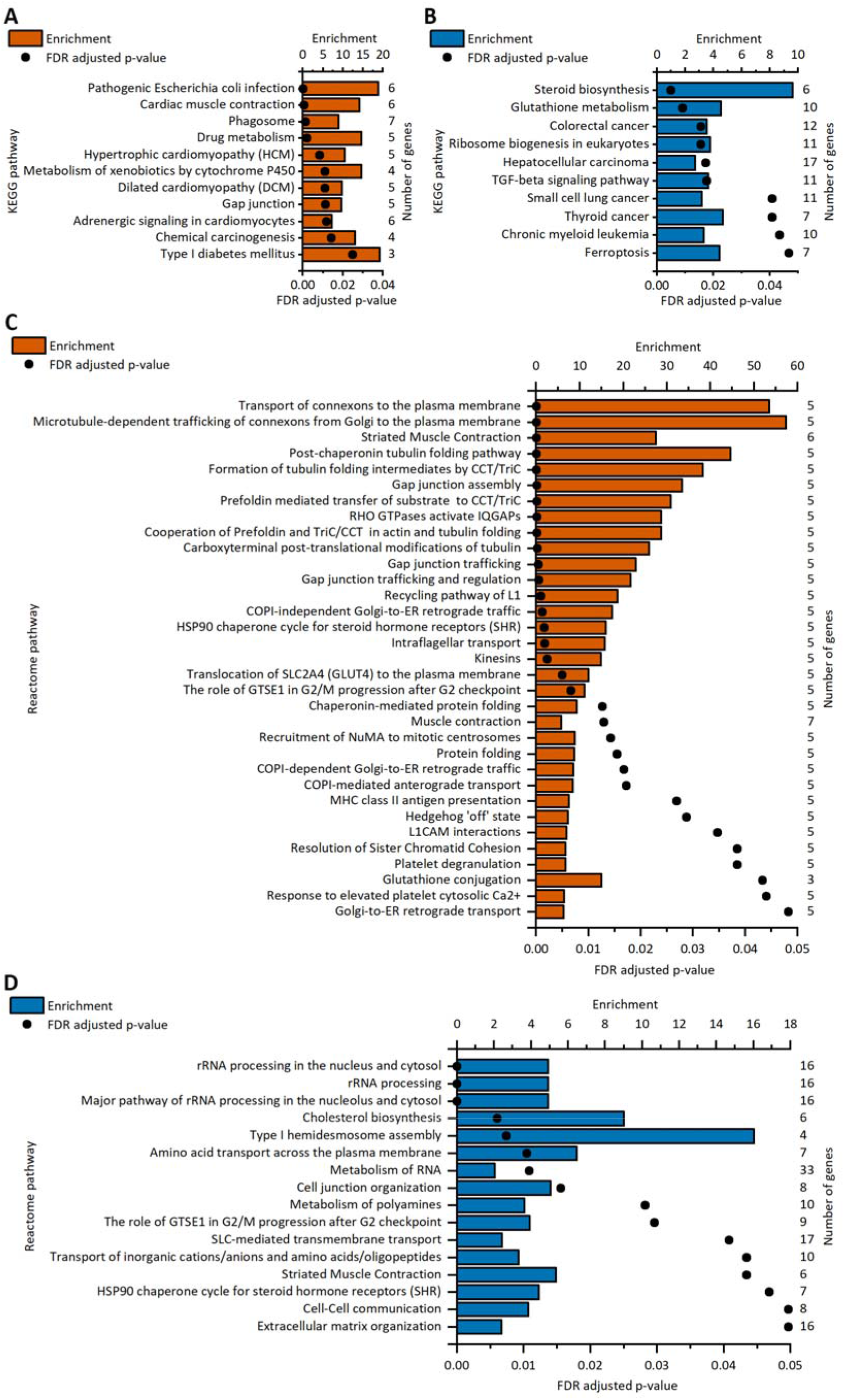
Enriched pathways in upregulated genes after cyclic stretch in human induced pluripotent stem cell-derived cardiomyocytes (hiPSC-CMs) and neonatal rat ventricular myocytes (NRVMs). A-B, KEGG pathway analyses of upregulated genes of stretched hiPSC-CMs (A) and NRVMs (B). C-D, Reactome pathway analyses of upregulated genes of stretched hiPSC-CMs (C) and NRVMs (D). For each significantly enriched pathway term, enrichment values are presented as bars, and FDR-adjusted p values are presented as dots. The number of upregulated genes associated with each term is shown on the right. The upregulated genes in the NRVMs used for the analysis are from Rysä et al.^12^

### Enrichment of transcription factor targets sites in hiPSC-CMs and NRVMs

Several transcription factors (TFs) are associated with cardiomyocyte hypertrophy.^5,26^ Hence, we analyzed which TF target sites were enriched in response to stretch. In our analysis, 19 and 18 TF binding sites were enriched in upregulated genes of stretched hiPSC-CMs and NRVMs, respectively (Figures 8A and 8B). Most of the enriched binding sites were for serum response factor (SRF), which had 5 enriched binding sites in both cell types. SRF controls the expression of genes regulating the cytoskeleton during development and cardiac hypertrophy. ^27,28^ In addition, two binding sites for the transcription factor c-Jun, which is part of the AP-1 complex and is involved in cardiomyocyte hypertrophy and increased steroidogenic gene expression,^29,30^ were enriched in both hiPSC-CMs and NRVMs. Both cell types also had an enriched binding site for transcription factor E2-alpha (TCF3) and for nuclear factor erythroid-derived 2 (NFE2). In hiPSC-CMs, two binding sites for myocyte enhancer Factor 2A (MEF2A) were enriched. MEF2A is known to regulate multiple cardiac structural genes and to be activated by several hypertrophic signaling pathways.^31–33^ In NRVMs, two binding sites for both MYC proto-oncogene protein and heat shock transcription factor 1 (HSF1) were enriched. However, none of the transcription factors mentioned above were differentially expressed in stretched hiPSC-CMs. In NRVMs, two of these transcription factors were differentially expressed.^12^ A gene coding for transcription factor c-jun was upregulated after 48 h of stretch and a gene coding for myc was upregulated both after 24 h and 48 h of stretch.

**Figure 8.**
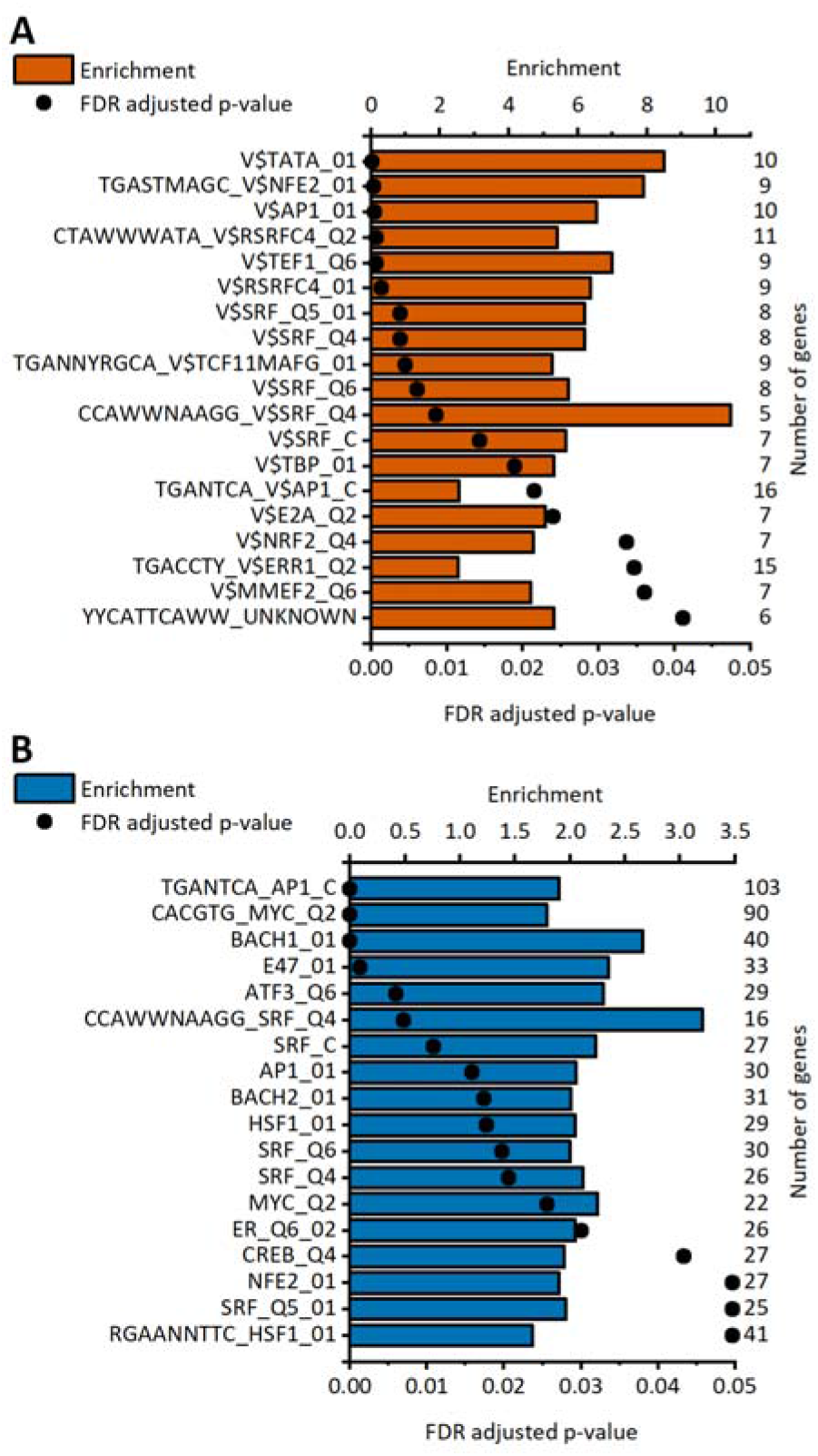
Enriched transcription factor target sites in the upregulated genes of the stretched human induced pluripotent stem cell-derived cardiomyocytes (hiPSC-CMs; A) and neonatal rat ventricular myocytes (NRVMs; B). Enrichment values for each significantly enriched target site are presented as bars, and FDR-adjusted p values are presented as dots. The number of upregulated genes associated with each target site is shown on the right. The upregulated genes in NRVMs used for the analysis are from Rysä et al.^12^

### Differentially expressed lncRNAs

Of the 7,818 long noncoding RNAs (lncRNAs) expressed in our samples, only one lncRNA was differentially expressed after 24 h of stretch: *LINC00648* was downregulated by 50% (p=0.011) relative to the unstretched control. *LINC00648* also showed downregulation by 30% in hESC-CMs after a 48-h stretch.^13^ In hiPSC-CMs, after 48 h of stretch, one lncRNA (*LINC00702*) was upregulated, while five lncRNAs (*AUXG01000058*.*1, AZIN1-AS1, LAMTOR5-AS1, LINC01341, PTPRG-AS1*) were downregulated. There is no previous report of these lncRNAs being involved in cardiomyocyte hypertrophy. Predicted putative interaction partners of the differentially expressed lncRNAs are shown in Table S4. None of these interaction partners was differentially expressed (>1.5-fold) in response to stretch. However, an interaction partner of *AZIN1-AS1*, a gene encoding Egl-9 family hypoxia inducible factor 3 (*EGLN3)*, was upregulated 1.48-fold (p=0.0454). EGLN3 is a prolyl hydroxylase that is activated by hypoxia and may regulate cardiomyocyte apoptosis.^34^

### The effects of p38 MAPK, MEK1/2 and PKC inhibition on the stretch response

To gain further insight into the signaling routes mediating the stretch response in hiPSC-CMs, we investigated the involvement of p38 MAPK, MEK1/2 and PKC signaling pathways in the hiPSC-CM stretch response. We treated the cells with pharmacological inhibitors of these signaling molecules, stretched the cells for 24 h and analyzed mRNA expression of well-established stretch biomarkers *NPPA* and *NPPB*, as well as selected differentially expressed genes identified in this study, namely *ACTA1, GAL, CSRP3, SLC16A9*, and *LINC00648*. The genes *NPPA, CSRP3* and *SLC16A9*, which were dysregulated only modestly in response to 24-h stretch in RNAseq analysis, were not significantly affected by stretch in this experiment (Figure S6). In contrast, genes showing more pronounced changes in RNAseq analysis, were significantly dysregulated. Increased *NPPB* (5.6-fold; p=0.008), *ACTA1* (8.0-fold, p=0.008) and *GAL* (10-fold, p=0.008) expression and decreased LINC00648 expression (60%, p=0.008) was detected in stretched control cells compared to unstretched control cells (Figure 9).

**Figure 9.**
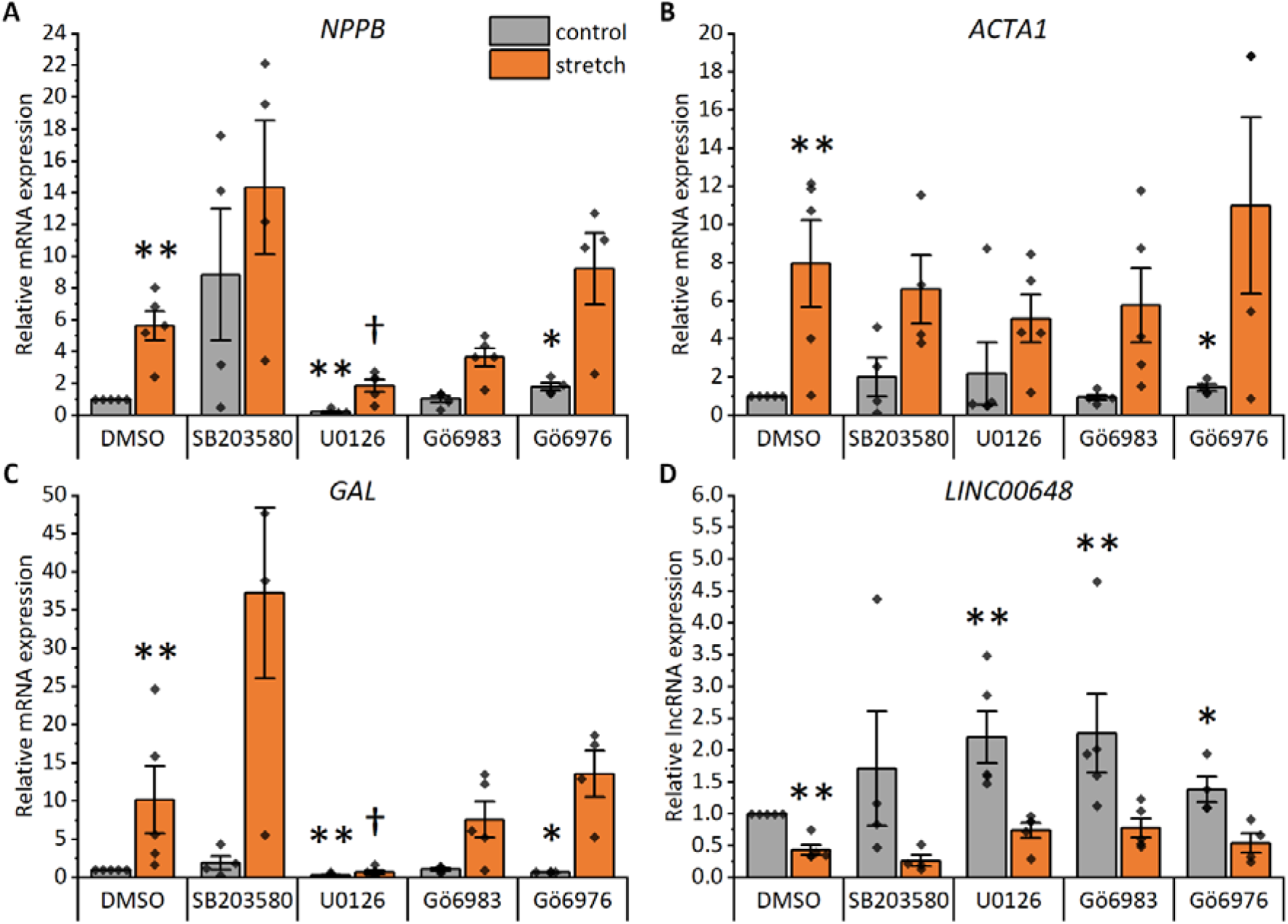
Effects of p38 mitogen-activated protein kinase (p38 MAPK), mitogen-activated protein kinase kinase 1/2 (MEK1/2) and protein kinase C (PKC) inhibitors on stretch-induced hypertrophic gene expression in human pluripotent stem cell-derived cardiomyocytes (hiPSC-CMs). The following inhibitors were utilized: SB203580 at 10 µM to inhibit p38 MAPK, U0126 at 10 µM to inhibit MEK1/2, Gö6983 at 1 µM to inhibit all PKC isoforms and Gö6976 at 1 µM to inhibit classical PKC isoforms. Natriuretic peptide B (*NPPB*; A), skeletal muscle alpha actin 1 (*ACTA1*; B), galanin (*GAL*; C) and long intergenic non-protein coding RNA 648 (*LINC00648*; D) RNA expression was measured with qRT–PCR after a 24-h cyclic mechanical stretch. The results are presented as fold change relative to the unstretched control. The data are shown as mean ± standard error of the mean, and values from individual experiments are presented as dots (n=5, except for SB203580 and Gö6976 n=4; where n represents biological replicates of cells from individual differentiations). *p<0.05, **p<0.01 vs. unstretched DMSO, † p<0.05 vs. stretched DMSO, Mann–Whitney U test.

Two of the inhibitors had significant effects on *NPPB* expression (Figure 9A). The MEK1/2 inhibitor U0126 at a concentration of 10 µM decreased the *NPPB* expression both in unstretched and stretched hiPSC-CMs compared to control (DMSO). However, it was not able to block the stretch response completely. In contrast, the inhibitor of classical PKC isoforms Gö6976 at 1 µM increased basal *NPPB* expression, but had no effect on stretch-induced *NPPB* expression. The p38 MAPK inhibitor SB203580 at 10 µM and the pan-PKC inhibitor Gö6983 at 1 µM had no effect on *NPPB* expression.

The expression of *ACTA1* was only affected by the classical PKC inhibitor Gö6976, which slightly increased the basal expression (Figure 9B). None of the inhibitors affected the stretch-induced upregulation of *ACTA1*. However, U0126 showed tendency towards stretch-reversing effect on *ACTA1* expression.

Both U0126 and Gö6976 decreased the baseline *GAL* expression (Figure 9C). However, only U0126 was able to inhibit the stretch-induced *GAL* expression. Pan-PKC inhibitor Gö6983 showed no effect on GAL expression, while p38 MAPK inhibitor SB203580 showed high variability but no clear influence on the *GAL* expression.

The MEK1/2 inhibitor and both PKC inhibitors increased the *LINC00648* expression at baseline (Figure 9D). However, none of the inhibitors affected the stretch-induced decrease in *LINC00648* expression, although U0126 and Gö6983 showed a tendency towards stretch-reversing effect.

Overall, since none of the inhibitors was able to block the stretch response completely, none of the pathways investigated alone is fully responsible for the stretch response of these four genes. The results however suggest involvement of MEK1/2 and PKC in mediating the hiPSC-CM stretch response.

## Discussion

Prolonged mechanical load leads to maladaptive changes in the heart, including cardiomyocyte hypertrophy and left ventricular hypertrophy, which are major causes of heart failure.^5^ Understanding the molecular mechanisms that underlie the development of left ventricular hypertrophy is essential for finding new treatments for heart failure. Identification of genes and pathways involved mechanical stretch response of cardiomyocytes is therefore of great interest.

The optimal *in vitro* model of cardiomyocyte hypertrophy would utilize adult human cardiomyocytes. However, they are difficult to obtain and culture long-term, therefore the most commonly used *in vitro* hypertrophy models employ NRVMs. To study this phenomenon in human cells and reduce the use of experimental animals, we used hiPSC-CMs, which are increasingly used as *in vitro* models in disease modeling and drug development.^35–37^ To our knowledge, the transcriptional responses of hiPSC-CMs to cyclic mechanical stretch have not been characterized before. We validated the model by measuring the expression of well-established mechanical stress-responsive genes, *NPPA* and *NPPB*,^25^ and showed that cyclic mechanical stretch leads to *NPPA* and *NPPB* gene expression responses comparable to those of NRVMs.^12,38^

We then performed RNAseq to identify additional gene expression changes involved in the hypertrophic response. Surprisingly, the number of differentially expressed genes in the stretched hiPSC-CMs was drastically lower than in the stretched NRVMs or hESC-CMs.^12,13^ In the NRVM study,^12^ the species difference and the different methods (microarray in the NRVM study and RNAseq in the present hiPSC-CM analysis) may partly explain the difference. Furthermore, the hiPSC-CM cultures used in the current analysis were ≥ 95% pure cardiomyocytes, while NRVMs isolated from the heart usually contain other cardiac cell types, such as fibroblasts and endothelial cells, despite cardiomyocyte enrichment with preplating. In agreement with this, a broad range of gene expression changes was observed in NRVMs, including several changes associated with cardiac fibroblast activation such as upregulation of α-smooth muscle actin (*ACTA2*), a hallmark of fibroblast activation, and overrepresentation of GO terms of extracellular matrix and collagen-containing extracellular matrix. Similar changes in gene expression were not detected in stretched hiPSC-CM, suggesting that pure cardiomyocyte cultures are more suitable to study the changes occurring specifically in cardiomyocytes. In the hESC-CM study,^13^ >98% pure cardiomyocytes were used, but the magnitude and frequency of the stretch applied were different. Moreover, the age and maturation level of the cardiomyocytes were not described in the hESC-CM study, and these differences may influence the comparison of the two human cardiomyocyte models. Although transcriptionally hESC-CMs and hiPSC-CMs are very similar,^39^ their responses to stretch were different. As NRVMs, hESC-CMs also seem to respond to stretching by inducing a broad range of gene expression changes, while in hiPSC-CMs, differentially expressed genes are more defined. Only one gene, *TUBB2B*, coding for tubulin beta-2A chain, a constituent of microtubules, was upregulated in all cardiomyocyte types. In hiPSC-CMs, other forms of alpha- and beta-tubulins were also upregulated. The increase in the expression of microtubules is strongly associated with cardiac hypertrophy; thus, this change was anticipated.^40^ In contrast, one gene, *ZNF519*, coding for zinc finger protein 519, was downregulated in all cell models. *ZNF519* has not been characterized in cardiomyocytes, and its potential role in the development of cardiomyocyte hypertrophy remains to be established.

In response to stretch, two central changes occur in cardiomyocytes: (1) several genes normally expressed only in embryonic or fetal hearts are reactivated, and (2) the expression of sarcomeric and other constitutive proteins is increased.^41,42^ Here, we showed that these changes occur also in hiPSC-CMs in response to stretching. The upregulation of contractile proteins was also reflected in the GO enrichment analysis, where most enriched processes were associated with actin-myosin filament sliding and muscle contraction.

Although the differentially expressed genes and their numbers varied in the compared cardiomyocyte models, some similarities in the enriched processes and pathways were discovered. Regulation of cell death and sterol biosynthesis were enriched in the upregulated genes in all cell types. Apoptosis has previously been linked to hypertrophy in multiple studies both in rodents and in humans.^43–46^ Although the upregulation of genes associated with steroid biosynthesis has been reported in previous studies,^12,13^ its role in cardiomyocyte hypertrophy has not been characterized. Increased steroid synthesis might be needed for the growth of cardiomyocytes or may be associated with changes in energy metabolism. Steroid biosynthesis is downregulated in the neonatal mouse heart within the first nine days of postnatal life, during which the heart loses its regenerative capacity.^47^ Hence, it can be speculated that increased steroid synthesis is a part of the fetal program that is reactivated in response to stress.

Several genes associated with both apoptosis and cardiomyocyte hypertrophy, such as *CRYAB, ENO1* and *GSTO1*, were among the upregulated genes in hiPSC-CMs.^48–55^ *ENO1*, which codes for the glycolytic enzyme α-enolase and is normally highly expressed in embryonic and fetal heart but only weakly in adult heart, has shown to increase during hypertrophy in animal models.^56–58^ This is in line with previous evidence of a metabolic switch from fatty acid to glycolysis during pathological hypertrophy.^59^ Furthermore, one study has shown compensatory increase in α-enolase expression to protect cardiomyocytes from hypertrophy.^56^ Interestingly, after a 48-h stretch, the most upregulated genes were galanin and GMAP prepropeptide coding gene *GAL*. Galanin is expressed principally in the nervous system and in some peripheral organs, but no expression in cardiomyocytes has been reported.^60^ However, its receptors are expressed in various cell types, including cardiomyocytes, and it has been suggested to be cardioprotective.^60–64^

All cardiomyocyte types included in the present comparisons, hiPSC-CMs, hESC-CMs and NRVMs, are considered relatively immature and do not fully correspond to adult cardiomyocytes in terms of their sarcomere structure, metabolism, or electrophysiological properties.^8^ However, based on our comparison, stretched hiPSC-CMs were the only cell model in which biological processes of muscle contraction and actin-myosin filament sliding were enriched among the upregulated genes. In view of *in vivo* cardiac overload, these are the most important processes to enhance in order to preserve cardiac pump function. However, these changes could also imply maturation of hiPSC-CMs, but this is unlikely because they were accompanied by upregulation of the fetal gene program and apoptosis-associated genes. Moreover, hiPSC-CMs were the only cells in which upregulated genes had enrichment of pathways for hypertrophic cardiomyopathy. On the contrary, these pathways were enriched among downregulated genes of stretched hESC-CMs.^13^ Taken together, hiPSC-CMs show distinct hypertrophic changes in gene expression at the levels of individual genes and biological processes, indicating that cyclic stretching of hiPSC-CMs is an advantageous *in vitro* model for studying mechanically induced cardiomyocyte hypertrophy.

When comparing the gene expression changes of stretched hiPSC-CMs to another widely utilized hypertrophic stimulation, ET-1, we identify only few similarities. Johansson et al. performed RNAseq for 24-h, 48-h and 72-h ET-1-treated hiPSC-CMs.^65^ In that study, 696, 152 and 163 genes were dysregulated after 24-h, 48-h and 72-h ET-1 treatment, respectively. The 23 upregulated genes, which were common to our stretch model, included for instance fetal genes (*ACTA1, NPPB, TAGLN*) and structural protein coding genes (*ACTC1, KRT8, KRT18, TUBB2A, TUBB2B, TUBB6*), while the three common downregulated genes were *DLG2, PLCG2* and *SLC35E2A*. In addition, Aggarwal et al. performed mRNA profiling for 18-h ET-1-treated hiPSC-CMs.^66^ In that study, 235 and 290 genes were up- and downregulated, respectively, in response to ET-1. The 15 upregulated genes, which were common to our stretch model, included again fetal genes (*ACTA1, NPPB, TAGLN*) and structural protein coding genes (*ACTN1, KRT8, KRT18, TUBA4A, TUBB2A*), while the seven common downregulated genes included for example the non-coding RNA *LINC00648* and *SLC16A9*, a gene coding a transporter protein. This suggests that, in hiPSC-CMs, neurohumoral and mechanical hypertrophic stimuli induce largely different gene expression responses.

Recently, Nicin et al. profiled gene expression changes in cardiomyocytes of human hypertrophic heart.^67^ They found 527 upregulated and 2,775 downregulated genes in cardiomyocytes of aortic valve stenosis (AS) patients compared to healthy hearts. Nine of the upregulated genes (*ACTA1, ACTN1, CYB5R1, ENO3, MRPL41, NES, TNNC1, TNNI3, TPM2*) and three of the downregulated genes (*AZIN1-AS1, DLG2, SDK1*) were common to our hiPSC-CM model. However, substantial differences in the number of dysregulated genes were observed. Considering the stage of the hypertrophy (early stage in our model and late stage in AS), these considerable differences in gene expression were anticipated. For reasonable comparison of genetic changes of hiPSC-CMs to late-stage hypertrophic heart, prolonged stretching of hiPSC-CMs is required. Given that hiPSC-CMs, unlike NRVMs, can be cultured for long periods (even months or years), prolonged stretching of hiPSC-CMs would be feasible and interesting. Nevertheless, a 2-dimensional monotypic cell culture lacking neurohumoral stimuli and heterotypic cell communication would not be expected to yield identical gene expression changes as observed *in vivo*.

Finally, we applied our model to study signaling pathways associated with cardiomyocyte hypertrophy. Previous studies have shown that mechanical stretch induces activation of multiple kinases, including PKC, MEK1/2-ERK1/2 and p38 MAPK both *in vitro* in NRVMs and *in vivo* in rats.^12,68–73^ Although these kinases are widely studied in animal models, their role in human cardiomyocytes is poorly known. Földes et al. showed that inhibition of p38 MAPK abolished both phenylephrine- and stretch-induced increase in hESC-CM size.^74^ In addition, MEK1/2 inhibition slightly inhibited phenylephrine-induced hypertrophy of hESC-CMs. Moreover, our previous study demonstrated that MEK1/2 mediates ET-1-induced proBNP and *NPPB* expression in hiPSC-CMs.^10^ These results suggest that p38 MAPK and MEK1/2-ERK1/2 pathways are related to the hypertrophic response also in human cardiomyocytes. To our knowledge, the role of PKC in stretch-induced hypertrophy has not been studied human cardiomyocytes before. However, we have previously shown that activation of all PKC isoforms or inhibition of classical PKC isoforms induce pro-hypertrophic changes in hiPSC-CMs.^10^ Therefore, we were interested to study the genetic response of hiPSC-CMs to inhibition of these kinases in combination with cyclic mechanical stretch. Interestingly, MEK1/2 inhibition blocked *GAL* expression almost completely in both unstretched and stretched hiPSC-CMs. The inhibitory effect of MEK inhibition on *GAL* expression has previously been shown in neuronal cells, but not in cardiomyocytes.^75^ Hence, it seems that MEK1/2-ERK1/2 is a common pathway for *GAL* expression regardless of the cell type. In addition, MEK1/2 inhibition markedly decreased stretch-induced *NPPB* expression, but could not fully block it, unlike in ET-1-induced hiPSC-CM hypertrophy.^10^ In hiPSC-CMs, ET-1 and mechanical stimulation thus seem to induce BNP expression through different signaling pathways, which is in line with the differences in gene expression profiling data from stretched and ET-1-treated hiPSC-CMs.^66^ Overall, these results suggest that the stretch-induced gene expression changes are mediated through different signaling pathways, and that at least for some of the genes studied here the stretch-response is regulated via multiple intracellular signaling pathways.

We recognize that the hypertrophy model of stretched hiPSC-CMs has its limitations. First, as mentioned before, the hiPSC-CMs are not perfectly resembling adult cardiomyocytes and they are relatively immature in many regards.^8^ However, our results suggest that the gene expression response of these cells resembles adult human cardiomyocytes better than that of NRVMs. Second, hiPSC-CMs are cultured in isolation of other cardiac cell types. Hence, they lack the heterotypic cell-cell interactions that are present in normal cardiac environment. Single-cell sequencing of co-cultured cells could be performed to study the hypertrophic responses of cardiomyocytes in heterocellular environment. Third, our model reflects the onset of cardiomyocyte hypertrophy, and is thus not comparable to cardiomyocyte hypertrophy of late-stage hypertrophic heart. The model could however be further utilized to study long-term effects. Finally, the study was conducted using only one healthy hiPSC line. However, only minimal differences have been demonstrated in cardiomyocytes produced by the differentiation protocol used here.^14^ Therefore, similar stretch responses could also be predicted in other hiPSC lines with no known genetic mutations affecting cardiomyocyte function.

In conclusion, in the present study, we showed that mechanical stretching of hiPSC-CMs is a relevant *in vitro* model for studying human cardiomyocyte hypertrophy. We elucidated stretch-induced transcriptional changes and identified biological processes and pathways associated with these gene expression changes. The changes, including activation of the fetal gene program and upregulation of constitutive protein coding genes, were characteristic of cardiomyocyte hypertrophy. Comparison to previous data of stretched NRVMs and hESC-CMs demonstrated that hiPSC-CMs revealed more defined changes in gene expression and that differentially expressed genes were restricted to cardiac and hypertrophy-related genes. In addition, we identified several differentially expressed genes with no or weak previous association with cardiomyocyte hypertrophy. These results can be utilized to further elucidate hypertrophic signaling pathways and to discover potential pharmacological targets and biomarkers of cardiomyocyte hypertrophy.

## Acknowledgements

We thank Ms Annika Korvenpää and Ms Ángela Morujo for expert technical assistance. The RNAseq was provided by the Biomedicum Functional Genomics Unit at the Helsinki Institute of Life Science and Biocenter Finland at the University of Helsinki. We thank American Journal Experts (AJE) for English language editing.

## Sources of funding

This work was supported by the Academy of Finland [grants 321564, 328909], the Finnish Foundation for Cardiovascular Research, Sigrid Jusélius Foundation, University of Helsinki Research Funds, and Doctoral Programme in Drug Research.

## Disclosures

None.

## Supplemental Material

Figures S1-S6

Tables S1-S4

Datasets S1-S2

## Data Availability Statement

The RNAseq data are freely available at the NCBI Gene Expression Omnibus (http://www.ncbi.nlm.nih.gov/geo, accession number GSE186208). All other data are available from the corresponding author upon reasonable request.

## Notes

### Competing Interest Statement

The authors have declared no competing interest.

### Summary of Updates

The following section was added: Immunofluorescence staining, imaging and image analysis. The following sections were updated: Effects of mechanical stretch on natriuretic peptide expression in hiPSC-CMs; Enrichment of transcription factor targets sites in hiPSC-CMs and NRVMs; The effects of p38 MAPK, MEK1/2 and PKC inhibition on the stretch response; Discussion; Figures 1, 2, 3, and 9; Supplementary Figures S1, S2, and S6.

