## Supplemental Material for "Transcriptomics reveal stretched human pluripotent stem cell-derived cardiomyocytes as an advantageous hypertrophy model"

**Contents:**

Supplemental Figures S1-S6, p. 2-6

Supplemental Tables S1-S4, p. 7-10

Supplemental Dataset Legends, p. 10

### Supplemental Figures

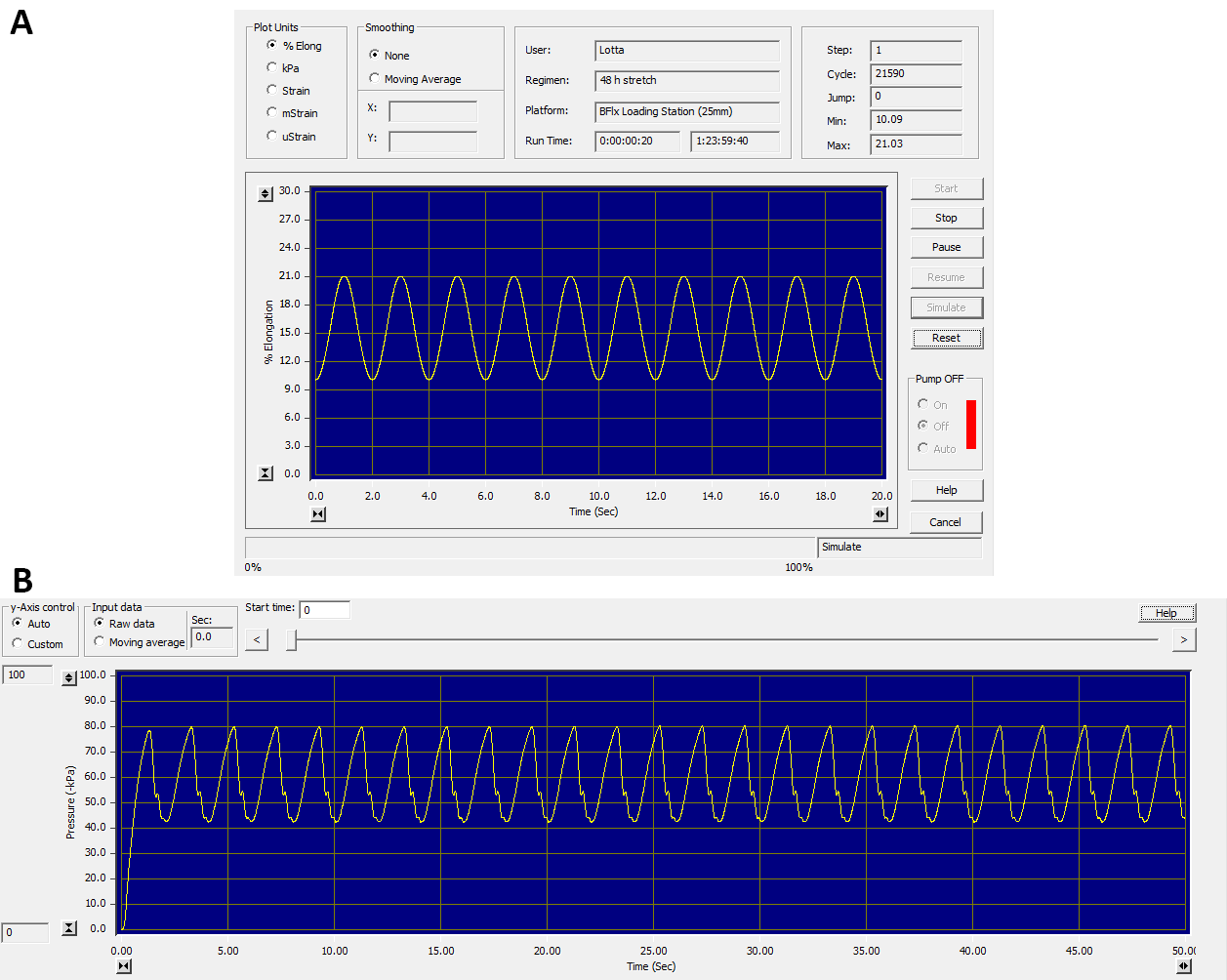

**Figure S1. A representative graph of waveform and degree of stretch.** **A**, A graph representing the start of a simulated regimen, where the membrane elongation (%) is plotted over time (sec). **B**, A graph representing the start of an actual regimen run, where the pressure (-kPa) is plotted over time (sec). Graphs are exported from the FX-5000™ software.

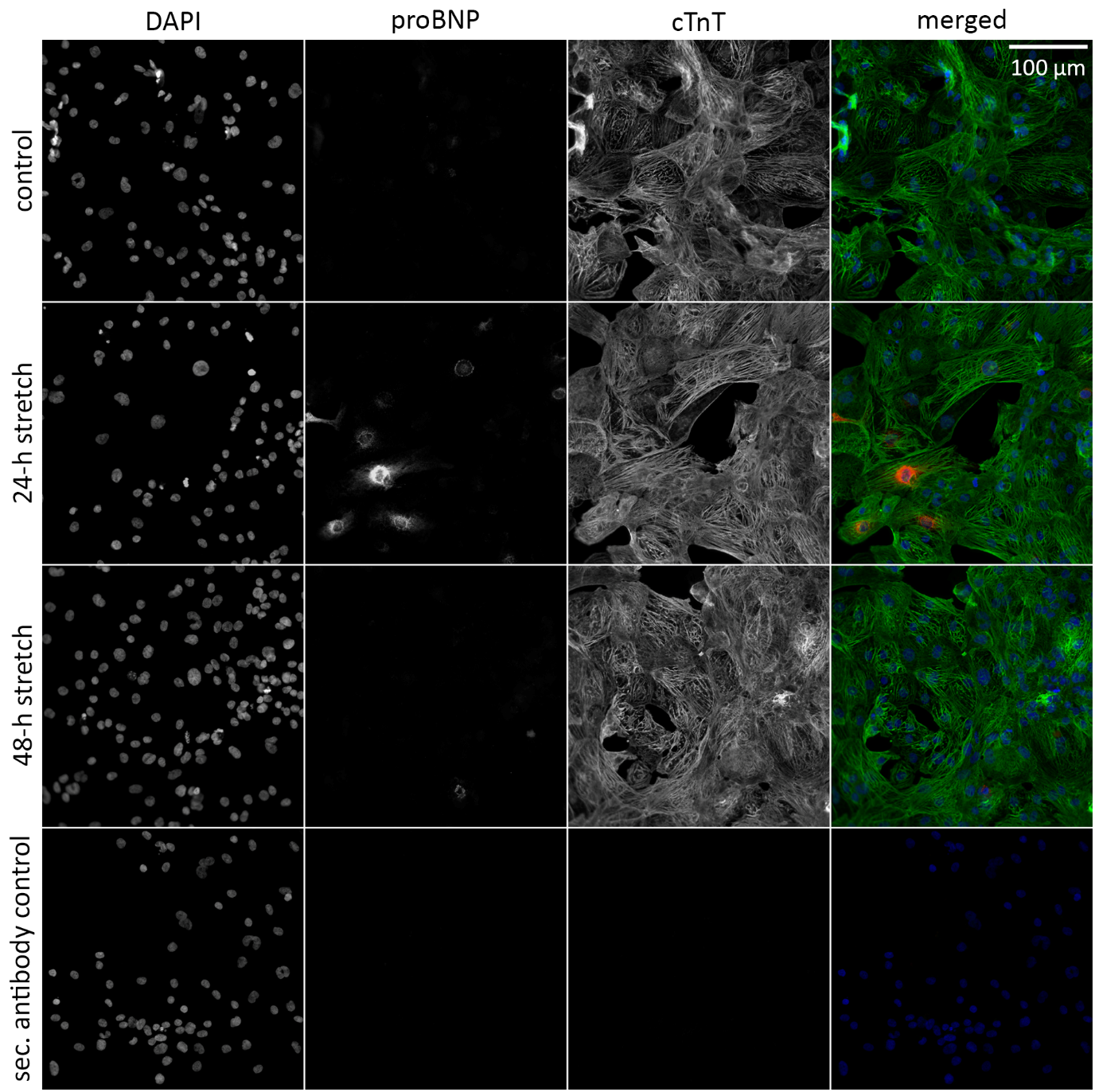

**Figure S2.** **Representative images of unstretched control, 24-h stretched, and 48-h stretched hiPSC-CMs.** Images are acquired with 40x water immersion objective. hiPSC-CMs were stained for DNA (DAPI; blue), pro-B-type natriuretic peptide (proBNP; red) and cardiac troponin T (cTnT; (green). Scale bar, 100 µm.

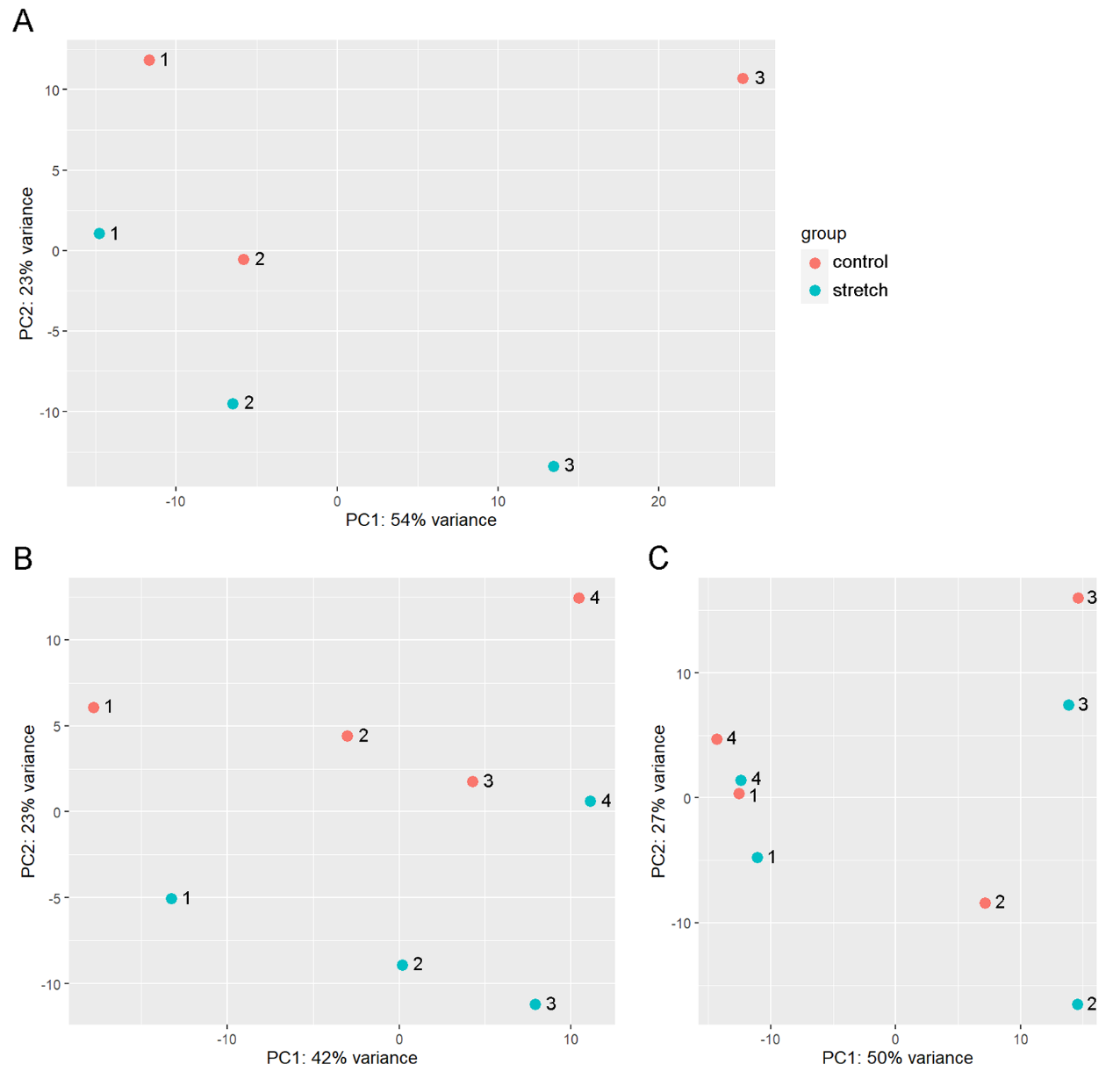

**Figure S3. Principal component (PC) analysis plots of the RNAseq results of the stretched and the unstretched control hiPSC-CMs at 24h (A), 48h (B) and 72h (C).** Each dot indicates a sample, and each sample pair is numbered (1-4). n=3 for 24h, n=4 for 48h and 72h.

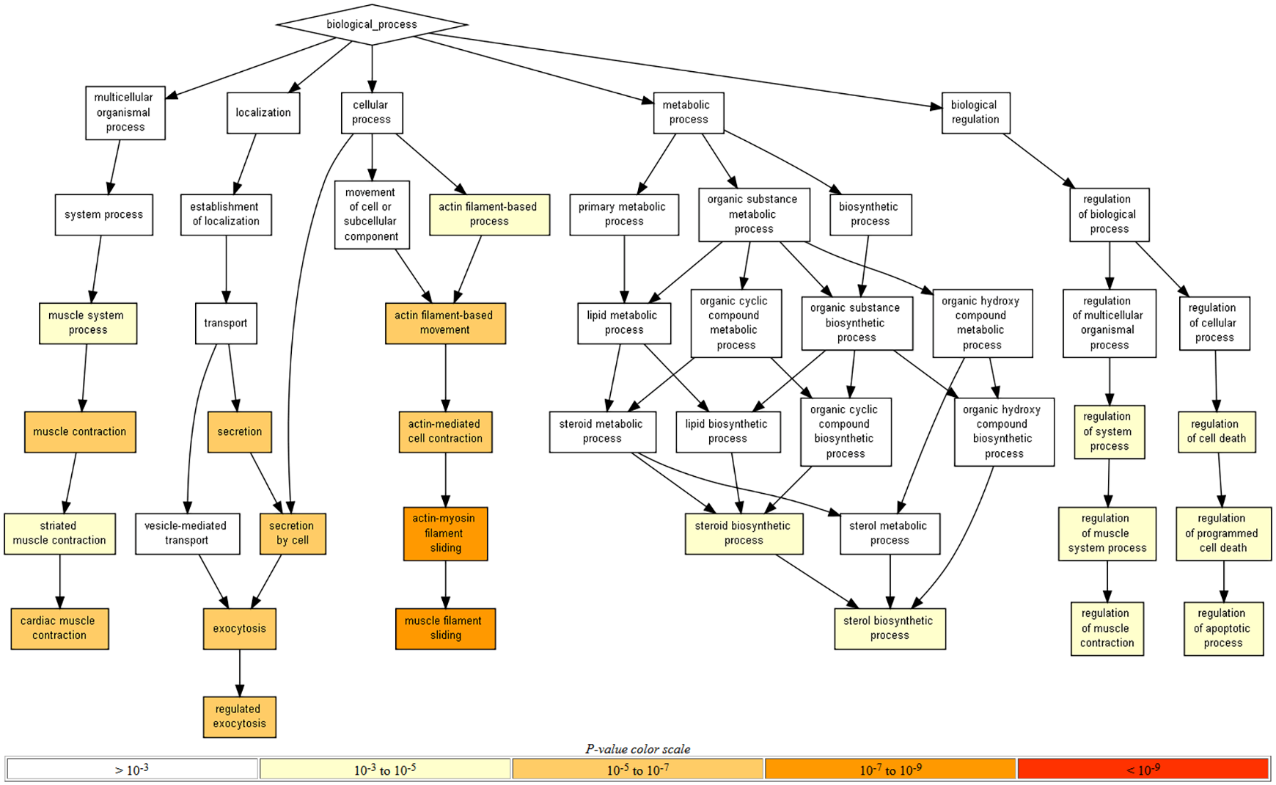

**Figure S4. Biological processes enriched in upregulated genes after 24h, 48h or 72h of cyclic stretch in hiPSC-CMs.** Gene Ontology (GO) enrichment analysis was performed with GOrilla. Directed graph of enriched processes is color-coded based on the significance of enrichment.

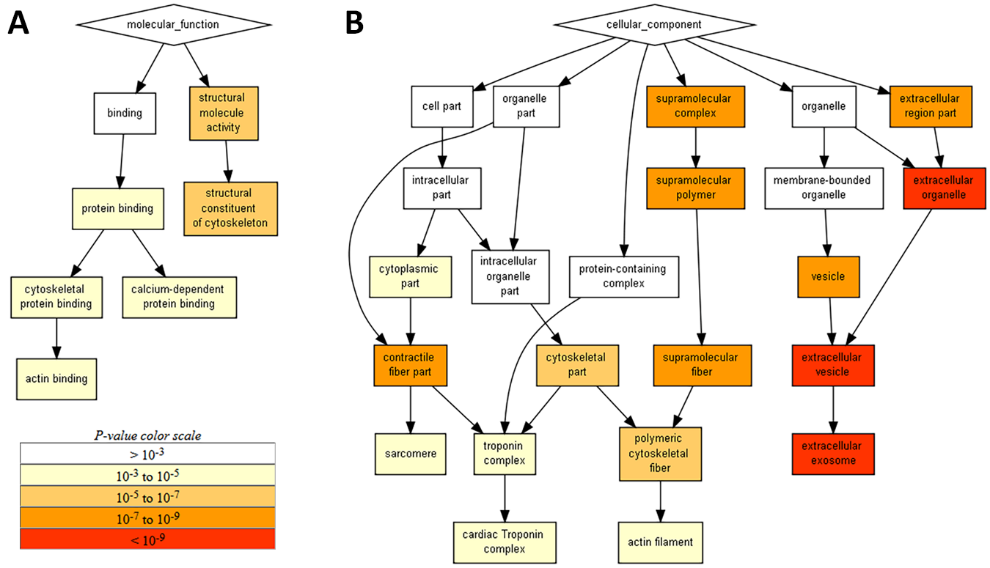

**Figure S5. Molecular functions (A) and cellular components (B) enriched in upregulated genes after 24-h, 48-h or 72-h cyclic stretch.**  Gene Ontology enrichment analysis was performed with GOrilla. Directed graphs of enriched molecular functions and cellular components are color-coded based on the significance of enrichment.

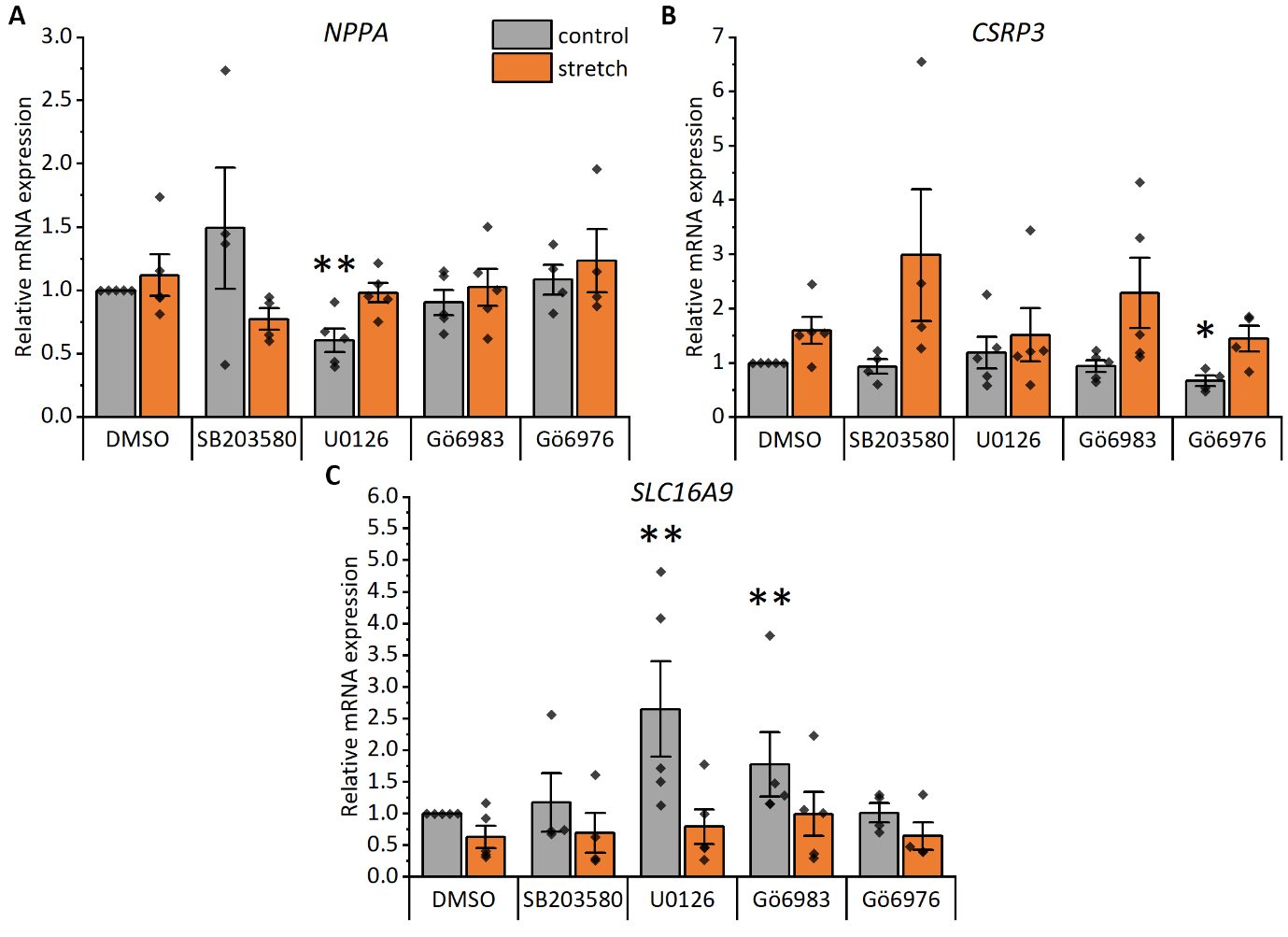

**Figure S6. Effects of p38 mitogen-activated protein kinase (p38 MAPK), mitogen-activated protein kinase kinase 1/2 (MEK1/2), and protein kinase C (PKC) inhibitors on NPPA, CSRP3 and SLC16A9 gene expression in hiPSC-CMs.** The following inhibitors were utilized: SB203580 at 10 µM to inhibit p38 MAPK, U0126 at 10 µM to inhibit MEK1/2, Gö6983 at 1 µM to inhibit all PKC isoforms, and Gö6976 at 1 µM to inhibit classical PKC isoforms. Natriuretic peptide A (NPPA; A), cysteine and glycine rich protein 3 (CSRP3; B), and solute carrier family 16 member 9 (SLC16A9; C) mRNA expression was measured with qRT–PCR after a 24-h cyclic mechanical stretch. The results are presented as fold change relative to the unstretched control. The data are shown as mean ± standard error of the mean, and values from individual experiments are presented as dots (n=5, except for SB203580 and Gö6976 n=4; where n represents biological replicates of cells from individual differentiations). *p<0.05, **p<0.01 vs. unstretched DMSO, † p<0.05 vs. stretched DMSO, Mann–Whitney U test.

### Supplemental Tables

**Table S1.** Used commercial TaqMan® Gene Expression Assays.

| **Target** | **Assay number** |
| --- | --- |
| *18S rRNA*, eukaryotic 18S ribosomal RNA | 4352930E |
| *ACTA1*, actin, alpha skeletal muscle | Hs00559403_m1 |
| *ACTC1*, actin, alpha cardiac muscle 1 | Hs01109515_m1 |
| *ACTN1*, alpha-actinin-1 | Hs00998100_m1 |
| *ACTB*, beta-actin | 4333762T |
| *CSRP3*, cysteine and glycine-rich protein 3 | Hs00185787_m1 |
| *GAL*, galanin | Hs00544355_m1 |
| *LINC00648*, long intergenic non-protein coding RNA 648 | Hs06598367_m1 |
| *NPPA*, natriuretic peptide A | Hs00383230_g1 |
| *NPPB*, natriuretic peptide B | Hs01057466_g1 |
| *PTPRG-AS1*, PTPRG antisense RNA 1 | Hs04970789_m1 |
| *SLC16A9*, solute carrier family 16 member 9 | Hs00415854_m1 |
| *TNNI3*, cardiac troponin I3 | Hs00165957_m1 |

**Table S2.** Protein families of the differentially regulated genes of the stretched hiPSC-CMs.

| **KEGG identifier** | **Protein family** | **Number of genes** |
| --- | --- | --- |
| **UPREGULATED** | | **77** |
| hsa01000 | Enzymes | 26 |
| hsa04147 | Exosome | 20 |
| hsa04812 | Cytoskeleton proteins | 15 |
| hsa03036 | Chromosome and associated proteins | 8 |
| hsa02000 | Transporters | 5 |
| hsa04131 | Membrane trafficking | 5 |
| hsa03019 | Messenger RNA biogenesis | 5 |
| hsa03029 | Mitochondrial biogenesis | 4 |
| hsa01009 | Protein phosphatases and associated proteins | 4 |
| hsa04090 | CD molecules | 3 |
| hsa03110 | Chaperones and folding catalysts | 3 |
| hsa03011 | Ribosome | 3 |
| hsa01002 | Peptidases and inhibitors | 3 |
| hsa04990 | Domain-containing proteins not elsewhere classified | 3 |
| hsa03000 | Transcription factors | 3 |
| hsa00537 | Glycosylphosphatidylinositol (GPI)-anchored proteins | 2 |
| hsa00199 | Cytochrome P450 | 1 |
| hsa01004 | Lipid biosynthesis proteins | 1 |
| hsa04050 | Cytokine receptors | 1 |
| hsa01007 | Amino acid related enzymes | 1 |
| hsa03016 | Transfer RNA biogenesis | 1 |
| hsa03041 | Spliceosome | 1 |
| hsa04040 | Ion channels | 1 |
| **DOWNREGULATED** | | **29** |
| hsa01000 | Enzymes | 6 |
| hsa03000 | Transcription factors | 6 |
| hsa04990 | Domain-containing proteins not elsewhere classified | 3 |
| hsa02000 | Transporters | 3 |
| hsa03036 | Chromosome and associated proteins | 3 |
| hsa03400 | DNA repair and recombination proteins | 2 |
| hsa04131 | Membrane trafficking | 2 |
| hsa01009 | Protein phosphatases and associated proteins | 2 |
| hsa01003 | Glycosyltransferases | 1 |
| hsa01001 | Protein kinases | 1 |
| hsa04147 | Exosome | 1 |
| hsa04040 | Ion channels | 1 |
| hsa01002 | Peptidases and inhibitors | 1 |
| hsa04515 | Cell adhesion molecules | 1 |

**Table S3.** Fold changes of the common differentially expressed genes of the stretched hiPSC-CMs, NRVMs (data from Rysä et al.^12^) and/or hESC-CMs (data from Ovchinnikova et al.^13^). Only statistically significant values are shown (p<0.05 for hiPSC-CMs and NRVMs and p<0.01 for hESC-CMs).

|  | **Fold change, stretch vs control** | | | | |
| --- | --- | --- | --- | --- | --- |
| **external gene name** | **24 h**  **hiPSC-CM** | **48 h**  **hiPSC-CM** | **24 h**  **NRVM** | **48 h**  **NRVM** | **48 h**  **hESC-CM** |
| *TUBB2B* | 2.7 | 2.6 | 2.3 | 2.0 | 2.2 |
| *CASQ1* | 1.8 | 1.6 | 2.6 | 1.8 |  |
| *TIMP1* | 1.7 | 1.7 | 1.5 | 1.6 |  |
| *ACAT2* | 1.8 | 1.7 |  | 1.8 | 1.8 |
| *CSRP3* | 2.0 | 1.7 |  | 1.5 |  |
| *TPM2* | 1.7 | 1.7 |  | 1.5 |  |
| *CNN1* | 2.1 |  | 4.0 | 7.4 | 1.9 |
| *MLLT11* | 1.7 |  | 2.7 | 3.7 | 1.6 |
| *NPPB* | 4.6 |  | 1.5 | 1.7 | 2.0 |
| *ACTN1* | 1.6 |  | 2.8 | 5.0 |  |
| *GADD45G* | 1.9 |  | 1.7 | 2.7 |  |
| *GPATCH4* | 1.6 |  | 1.7 | 1.5 |  |
| *TAGLN* | 3.6 |  | 2.0 | 1.7 |  |
| *TNFRSF12A* | 2.1 |  | 2.6 | 3.9 |  |
| *TUBB6* | 1.8 |  | 1.7 | 2.5 |  |
| *KRT18* | 1.9 |  | 1.5 |  |  |
| *SYPL2* | 1.5 |  | 1.7 |  |  |
| *PEA15* | 1.8 |  |  | 1.8 | 1.8 |
| *CYSTM1* | 1.6 |  |  | 1.5 |  |
| *NME1* | 1.7 |  |  | 1.8 |  |
| *PRSS23* | 2.1 |  |  | 1.5 |  |
| *TUBB2A* | 2.1 |  |  | 1.7 |  |
| *MASP1* | 2.0 |  | 0.6 | 0.5 | 1.6 |
| *CKB* | 2.2 |  |  |  | 1.7 |
| *TUBA4A* | 2.0 |  |  |  | 2.4 |
| *BOK* | 1.5 |  |  |  | 1.5 |
| *PTPRN* |  | 4.2 | 1.8 | 2.2 |  |
| *RCAN1* |  | 1.5 | 1.9 | 2.6 |  |
| *ENO3* |  | 1.6 |  | 0.5 | 1.6 |
| *DUSP13* |  | 2.8 |  |  | 2.4 |
| *CES1* |  | 2.0 |  | 0.4 |  |
| *GPCPD1* |  | 0.6 | 0.6 | 0.5 |  |
| *ZNF519* |  | 0.6 | 0.6 | 0.6 | 0.6 |
| *ADAM22* |  | 0.6 |  |  | 0.6 |
| *EGR1* |  | 0.5 |  |  | 0.5 |
| *EGR3* |  | 0.5 |  |  | 0.6 |
| *HELLS* |  | 0.6 |  |  | 0.5 |
| *HMGB2* |  | 0.7 |  |  | 0.6 |
| *MKI67* |  | 0.6 |  |  | 0.2 |
| *SKA3* |  | 0.5 |  |  | 0.4 |
| *TTYH2* |  | 0.6 |  |  | 0.5 |
| *DLG2* | 0.6 |  |  | 0.6 | 0.5 |
| *PLCG2* | 0.7 |  |  |  | 0.4 |
| *POLQ* | 0.7 |  |  |  | 0.5 |
| *SDK1* | 0.6 |  |  |  | 0.6 |
| *RAD51AP1* | 0.6 |  |  |  | 0.6 |
| *LINC00648* | 0.5 |  |  |  | 0.7 |

**Table S4**. The predicted interaction pairs for the differentially expressed lncRNAs of the stretched hiPSC-CMs.

| **LncRNA** | **Interaction Pair** |
| --- | --- |
| AZIN1-AS1 | PNN, SNRPN, CCDC32, KDM6A, FAM96B, SC5D, EGLN3, RHOH, AP003354.2, AC008124.1 |
| LAMTOR5-AS1 | KIF1B, MAPK6, ATP5S, RPL11, SLC1A5, GOSR2, SRD5A3, PPFIA1, U2AF1, ZNF621, PPIAP24, RNU6-179P, RNA5-8S5, RN7SL2, RN7SL703P, AL355488.1, U2AF1L5, AL139099.4, TDGF1, RNA28S5 |
| LINC00648 | SCAMP1, ARMC1, PRDM15, RPS27A, TMEM50A, AC002075.2, RPS6KC1, SYNPR-AS1 |
| LINC01341 | TKT |
| PTPRG-AS1 | SRSF7, AMPD2, RNA28S5 |

### Supplemental Dataset Legends

**Dataset S1.** Differential expression analysis of RNA sequencing.

**Dataset S2.** Gene Ontology analysis of the upregulated genes.
